# Regulation of chromatin transcription dynamics by DNA supercoiling

**DOI:** 10.1101/2023.11.06.565891

**Authors:** Sumitabha Brahmachari, Shubham Tripathi, José N Onuchic, Herbert Levine

## Abstract

Transcription has a mechanical component, as the translocation of the transcription machinery or RNA polymerase (RNAP) on DNA or chromatin is dynamically coupled to the chromatin torsion. This posits chromatin mechanics as a possible regulator of eukaryotic transcription, however, the modes and mechanisms of this regulation are elusive. Here, we first take a statistical mechanics approach to model the torsional response of topology-constrained chromatin. Our model recapitulates the experimentally observed weaker torsional rigidity of chromatin compared to bare DNA, and proposes structural transitions of nucleosomes into chirally distinct states as the driver of the contrasting torsional mechanics. Coupling chromatin mechanics with RNAP translocation in stochastic simulations, we reveal a complex interplay of DNA supercoiling and nucleosome dynamics in governing RNAP velocity. Nucleosomes play a dual role in controlling the transcription dynamics. The steric barrier aspect of nucleosomes in the gene body counteracts transcription via hindering RNAP motion, whereas the chiral transitions facilitate RNAP motion via driving a low restoring torque upon twisting the DNA. While nucleosomes with low dissociation rates are typically transcriptionally repressive, highly dynamic nucleosomes offer less of a steric barrier and enhance the transcription elongation dynamics of weakly transcribed genes via buffering DNA twist. We use the model to predict transcription-dependent levels of DNA supercoiling in segments of the budding yeast genome that are in accord with available experimental data. The model unveils a paradigm of DNA supercoiling-mediated interaction between genes and makes testable predictions that will guide experimental design.

---

Supercoiling of the genomic DNA is a ubiquitous feature of active transcription in both eukaryotes and prokaryotes. Translocation of the RNA Polymerase (RNAP), an active process generating RNA transcripts, overtwists the downstream DNA and undertwists the upstream DNA. First conceptualized in the twin-domain model more than three decades ago [1], the transcription-supercoiling interplay has come into renewed focus with recent experimental advances that allow tracking of individual transcribing RNAPs [2, 3] and genome-wide profiling of the DNA supercoiling [4, 5]. Transcription-generated supercoiling has been shown to speed up transcription elongation via collective RNAP behavior [3], influence gene burst kinetics [2, 6, 7], and impact the three-dimensional genome architecture [8–11].

Theoretical and computational models of the transcription-supercoiling interplay have been immensely useful in interpreting experimental observations and making testable predictions to guide experimental design [12–16]. These theoretical frameworks have to date focused on prokaryotic transcription and have accordingly incorporated the torsional response of bare DNA with varying levels of detail. However, the applicability of these models to eukaryotic transcription is unclear. This is because eukaryotic DNA predominantly resides in a nucleosome-wrapped state, termed chromatin, that is known to exhibit qualitatively different mechanics than bare DNA [17–19]. While experimental studies are increasingly probing the role of the supercoiling in eukaryotic transcription [5–8], there lacks a theoretical framework that quantitatively analyzes the transcription-supercoiling interplay in chromatin.

Nucleosomes can affect transcription in multiple ways, both chemical and mechanical. Chemically, histones, the constituent proteins of nucleosomes, serve as substrates for a variety of epigenetic modifications. These modifications can affect the recruitment of different components of the transcription machinery, as well as impact the three-dimensional genome architecture [20, 21]. Mechanically, nucleosomes can serve as steric barriers to RNAP recruitment and translocation [22]. Importantly, single-molecule assays have shown that nucleosomes alter the torsional response of bare DNA [17–19]. The observations suggest nucleosomes can act as torsional buffers, capable of absorbing or screening positive supercoiling. This effect has been phenomenologically incorporated into a model of the transcription-supercoiling interplay [15]. However, the absence of a quantitative model capable of predicting chromatin torsional response has held back a mechanistic treatment of supercoiling dynamics during eukaryotic transcription.

In this manuscript, we present a mechanistic framework to understand eukaryotic transcription and its regulation, a formulation that is inspired by our previous work on prokaryotes [14]. Within this framework, transcription initiation is simulated as a stochastic event where RNAPs are recruited at the transcription start site (TSS) at a rate that that sets the effective transcription initiation rate. Transcription elongation along topology-constrained (or net linking-number constrained) linear DNA, featuring translocation of the recruited RNAP and the associated transcription bubble, forces “arm wrestling” between the RNAP and the DNA. This is because the failure of a transcribing RNAP to rotate in congruence with the DNA groove results in an increased (reduced) DNA linking number density downstream (upstream). We simulate the DNA-twist-coupled translocation of RNAP via a set of dynamical equations that enforce torque balance between RNAP rotation and DNA twisting. This leads to increased rotation for less bulky RNAPs, while the DNA is twisted more when the RNAP bulk increases due to its attachment to larger mRNAs. The contest between RNAP rotation and DNA torsion is the same for prokaryotes and eukaryotes [14]. However, within our approach, eukaryotic transcription is distinct because of nucleosome binding.

Nucleosomes play a dual role within this framework. First, they may act as steric barriers to RNAPs, and second, they govern the torsional response of chromatin. We developed a statistical-mechanical model of chromatin that integrates nucleosome structural aspects with a twistable worm-like chain model of bare DNA. The model, in quantitative accord with experimental data [17, 18], shows a weak torsional stiffness of chromatin originating from transitions between the coexisting chiral states of nucleosomes (Fig. 1). These nucleosome states have varied writhe contributions to the DNA linking number and are structurally distinguishable based on the relative orientation of the two linker DNAs [17, 23]. We then use the model to analyze RNAP-induced DNA supercoiling in the context of chromatin and probe the kinetics of transcription elongation in eukaryotes (Fig. 2). While the steric hindrance aspect of nucleosomes counteracts transcription, we find that the weak torsional stiffness of chromatin facilitates transcription (Fig. 3). Finally, we use the framework to simulate transcription in various kilobase-scale segments of the yeast *S. cerevisiae* genome (Fig. 4). The model makes quantitative predictions regarding the supercoiling status of the segments, that are in agreement with the statistical trends in experimental data (Fig. 5 and Fig. 6). We also discuss how a perturbation in the expression level of one gene propagates over the segment, an effect driven by the altered levels of DNA supercoiling. Overall, our work argues that DNA supercoiling is an unavoidable and important aspect of actively transcribed eukaryotic DNA that has structural and functional consequences at multiple length scales.

**FIG. 1.**
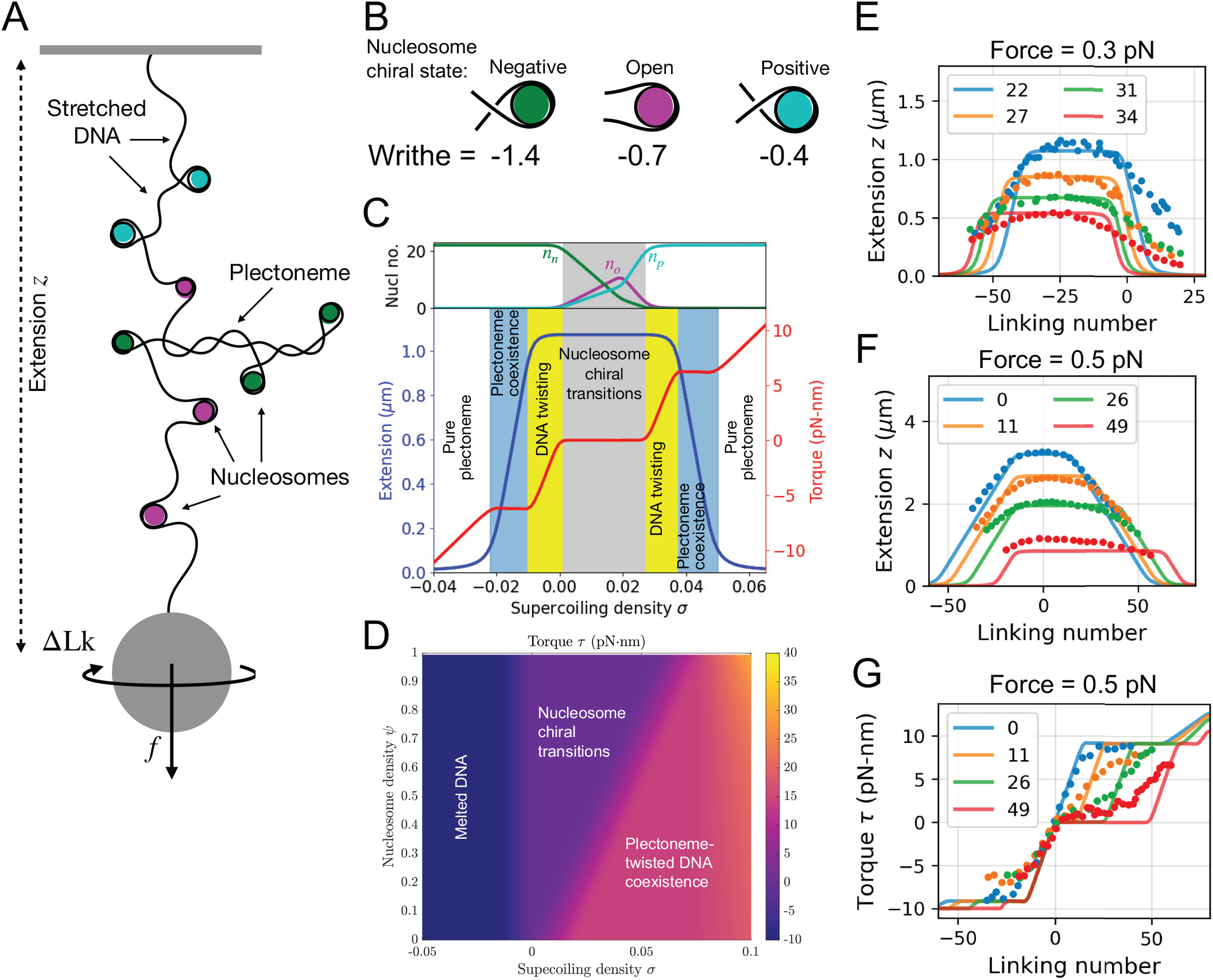
Torsional response of chromatin. **A** Schematic of the single-molecule tweezers setup commonly used to probe the torsional response of DNA / chromatin [17–19]. The two ends of the chromatin segment are torsionally constrained, such that one is fixed to the surface of a coverslip and the other to the surface of a bead. The excess linking number in the chromatin segment ∆Lk is controlled via the rotation of the bead. Additionally, the segment is put under a constant extensile force *f* . DNA in the chromatin fiber may wrap around nucleosomes, stretch under the external force, or buckle to form a plectoneme. **B** A DNA-bound nucleosome can exist in either a positive, open, or negative chiral state. These states store different amounts of linking number as writhe and they may interconvert via simple rotation about the dyad axis, thus changing their writhe contribution to the linking number of the DNA segment. **C** Chromatin fiber extension (blue; left vertical axis) and torque (red; right vertical axis) as a function of the chromatin supercoiling density *σ* shown for a DNA segment of length 8.2 kb containing *N* = 22 nucleosomes, under a force of 0.3 pN, and using a references state writhe Wr_ref_ = *N* Wr_*n*_. The distribution of nucleosomes among the different chiral states is shown in the top panel. Note that injecting positive supercoils into the chromatin fiber leads to a flat regime in extension and a low torque valley, which is due to the coexistence of nucleosome chiral transitions. Beyond this valley, nucleosomes are unable to accommodate or buffer DNA twists leading to a chromatin response similar to that of bare DNA. The width of the low-torque valley increases with the number of nucleosomes, as a higher number of chiral transitions are able to buffer more DNA twists. **D** Chromatin restoring torque for various supercoiling densities *σ* and nucleosome densities *ψ* at force *f* = 1.0 pN. Here, *ψ* = 0 corresponds to bare DNA and *ψ* = 1.0 corresponds to a chromatin fiber completely coated with nucleosomes with no free DNA. This *σ*-*ψ* dependence of the restoring torque was used for all the simulations of chromatin transcription in Fig. 2-6. **E** Chromatin extension *z* versus excess linking number ∆Lk under *f* = 0.3 pN for a 8.2 kb DNA segment. The different colors correspond to different numbers of nucleosomes *N* as shown in the legend. Solid curves correspond to predictions from our model using Wr_ref_ = 0, while the dots indicate the experimental observations from Bancaud *et al*. [17]. **F** Same as (E) for a 11.8 kb DNA segment under a 0.5 pN force, where *N* = 0 (blue) represents bare DNA. **G** DNA restoring torque corresponding to the setup in **F**. Solid curves in **F** and **G** correspond to predictions from our model using Wr_ref_ = *N* Wr_*n*_, while the dots indicate the experimental observations from Le *et al*. [18].

## I. RESULTS

### A. A statistical mechanical model incorporating chromatin topology, mechanics, and nucleosome chiral transitions

We model chromatin as a string of nucleosomes wherein each nucleosome is a structural unit that absorbs 60 nm (177 bp) of DNA (Fig. 1 A). This framework models nucleosome-driven DNA compaction and leads to a lower end-to-end extension of chromatin compared to bare DNA (Fig. 1 E). To probe the mechanics of chromatin we mimic the experimental setup of a single-molecule tweezers experiment, where chromatin is fixed at both ends, put under an extensile external force and a fixed rotation or linking number (Fig. 1A). The DNA in the chromatin can exist in stretched or plectonemic states and the nucleosomes can exhibit different chiral states (Fig. 1 B). While the stretched DNA state is stabilized by the extensile force and contributes to higher DNA extension, the plectonemic state arises when the applied DNA twist is large enough to buckle the DNA into a helically wrapped plectonemic configuration that stores linking number in the form of writhe, thereby absorbing DNA twist [24, 25].

The different nucleosome chiral states are defined by their configuration geometry (Fig. 1 B). Following previous studies [17, 26], we posit three topological, or chiral, states of nucleosomes: open, negative, and positive. These states store differing amounts of DNA writhe due to differences in the geometry of how the DNA linkers exit the nucleosome core. When the two DNA linkers do not overlap or cross each other, the nucleosome is in an “open” state (Fig. 1 B). Each nucleosome in the open state stores DNA writhe of Wr_*o*_ = − 0.7 which comes from the inner turn of the nucleosome. When the two linkers cross each other, there is an additional contribution to the total DNA writhe of the nucleosome. If the linker crossing has the same topological sense as the inner turn, the net writhe of the nucleosome is more negative, Wr_*n*_ = − 1.4, and we label it as the “negative” state (Fig. 1 B). In contrast, if the linker crossing has the opposite sense to the inner turn, we call it the “positive” state with a net DNA writhe of Wr_*p*_ = −0.4 (Fig. 1 B). These states can interconvert by rotations about the dyad axis. Such variations in nucleosome structure have been observed in cryo-electron microscopy studies [23]. These nucleosome states are otherwise considered identical, such as in terms of their DNA binding energy and DNA length absorption (see Fig. S2 and Fig. S3 for cases where this assumption is relaxed). Overall, a chromatin configuration with a fixed number of nucleosomes in open (*n*_*o*_), positive (*n*_*p*_), and negative (*n*_*n*_) states will have a total nucleosome writhe given by:

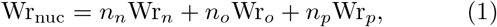

where *n*_*n*_ + *n*_*o*_ + *n*_*p*_ = *N* is the total number of nucleosomes. Note that the writhe values for these states were chosen based on previous studies [17, 19, 27].

We specify the overall chromatin state by simultaneously specifying details of the DNA and the nucleosome configurations. The DNA configuration is specified in the DNA fractions in a force-extended or stretched state and a plectonemically buckled state. The nucleosome configuration is specified by the number of nucleosomes in each of the three chirally distinct states (Fig. 1 A, B). We write the total free energy for a given chromatin state:

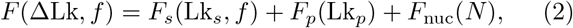

where the RHS terms are the contributions from stretched DNA, plectonemic DNA, and nucleosomal states, respectively (see Eq. S1 and Eq. S2). ∆Lk is the net change in the DNA linking number from a reference state; note that ∆Lk is often described in an intensive form as the supercoiling density *σ* ≡ ∆Lk*/*(*L*_0_*/h*) where *h*≈ 3.4 nm is the length of the DNA double-helix repeat and *L*_0_ is the total DNA length. Lk_*s*_ and Lk_*p*_ are contributions to the excess linking number from the stretched and plectonemic DNA states, respectively. The total free energy of the chromatin is minimized subject to the following linking number constraint:

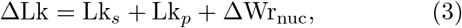

Here, ∆Wr_nuc_ ≡ Wr_nuc_ − Wr_ref_ is the deviation of nucleosomal writhe from the reference state Wr_ref_ . There are two possibilities for choosing Wr_ref_ . If a torsionally relaxed chromatin fiber is chosen as the reference, Wr_ref_ = *N* Wr_*n*_ is an appropriate choice. This assumes that in the reference state, all the nucleosomes are in a negative state. Single-molecule experiments where torsional constraints are added after nucleosome assembly [18], as well as the *in vivo* scenario, correspond to this choice of reference (Fig. 1 F, G). The other possibility is choosing the relaxed, bare DNA as the reference, *i*.*e*., Wr_ref_ = 0. Single-molecule experiments where nucleosomes are assembled on torsionally-constrained DNA [17] correspond to this choice. In such a scenario, the zero excess linking number state, which corresponds to relaxed bare DNA, has positively twisted DNA after nucleosome assembly [17] (Fig. 1 E). Note that either choice of Wr_ref_ ensures that ∆Lk = 0 in the reference state.

Finally, for a given excess linking number ∆Lk and extensile force *f*, we construct a partition function [25]:

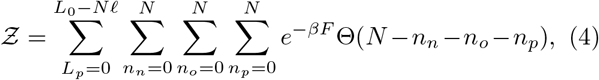

where Θ(*N*− *n*_*n*_− *n*_*o*_ − *n*_*p*_) = 1 if *n*_*n*_ + *n*_*o*_ + *n*_*p*_ = *N*, 0 otherwise. The torque follows:

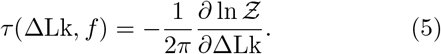

Other observables, like the end-to-end extension (Eq. S3) may be similarly obtained.

### B. Chromatin extension is less sensitive to excess twist than bare DNA due to nucleosome chiral transitions

When bare DNA is twisted, its extension initially remains unchanged since the force-extended state is stable in this regime. Beyond a threshold of excess twist, the DNA undergoes plectonemic buckling to compensate for the increasing DNA torque. Coexistence with the plectoneme state exhibits lower DNA extension as plectonemes do not contribute to extension [24, 25]. This behavior is shown by the *N* = 0 curve in Fig. 1 F . Modeling the chromatin twist response using the statistical mechanical model described above, we observe a qualitatively similar trend in the chromatin end-to-end extension as the bare DNA case (Fig. 1C, E, and F), *i*.*e*., a “hat”-shaped curve. There are, however, two key differences. First, untwisted chromatin has a lower extension than untwisted bare DNA. This is a consequence of the nucleosome-driven compaction of DNA: each nucleosome absorbs 60 nm of DNA that can no longer contribute to the end-to-end extension. Consistent with nucleosome-driven compaction, we observe that the end-to-end extension decreases with an increase in the number of nucleosomes (Fig. 1 E, F). Second, the top part of the “hat”-curve, *i*.*e*., the regime with flat end-to-end extension, is wider for chromatin as compared to bare DNA. Additionally, the stability of this regime, given by the width of the flat part, increases with an increase in the number of nucleosomes (Fig. 1 E, F).

When positive turns are injected into untwisted chromatin (the reference state with all nucleosomes in the negative state), the DNA does not twist in response. Instead, the nucleosomes undergo chiral transitions to a less negative state to accommodate the excess positive linking number. Negative nucleosomes first transition to the open states and then to the positive states (Fig. 1 C, top panel). As there is no buckling, the chromatin end-to-end extension does not change in this regime. Once all the nucleosomes have transitioned to a positive state, any additional linking number can only be accommodated by DNA twisting that finally leads to buckling, *i*.*e*., plectoneme formation. Increasing positive turns further increases the fraction of plectonemic DNA that does not contribute to extension leading to a steady decrease in end-to-end extension (Fig. 1 C, E, and F). Note that the open state is only transiently populated as shown in the top panel of Fig. 1 C. However, introducing a lower DNA binding energy for the open nucleosome state, as has been argued [17, 26, 27], leads to a stable open state at interim supercoiling densities (Fig. S2).

The stability of the unbuckled regime increases with the number of nucleosomes. This is because a larger number of chiral transitions allows for a DNA twist screening over a larger linking number range. As a result, the flat part of the “hat”-shaped regime increases with the number of nucleosomes (Fig. 1 E, F).

In contrast, when negative turns are injected into untwisted chromatin with all negative nucleosomes, the DNA immediately starts twisting since no nucleosome chiral transitions can accommodate negative supercoiling in this scenario. As more negative twists are injected, the DNA buckles and starts forming plectonemes with negative writhe (Fig. 1 C). Thus, chromatin’s response to negative twists is the same as for bare DNA. Note that at higher extensile forces (*f* ≈ 1.0 pN), the DNA may melt instead of forming negative plectonemes [24, 28].

Our predictions of the chromatin end-to-end extension in response to excess linking number injection are in agreement with the available data from two different single-molecule studies [17, 18] (Fig. 1 E, F). Note that we did not do any parameter fitting in our model. Combining the previously calibrated worm-like chain model for double-helix DNA [24] with the nucleosome parameters [17] in a consistent framework was enough to get the quantitative agreement with experimental data (Fig. 1 E-G).

### C. Nucleosome chiral transitions buffer DNA restoring torque

DNA twisting leads to a build-up of restoring DNA torque [24, 29]. For bare DNA, the restoring torque increases linearly with the excess linking number. However, when the DNA torque is above a critical value, it is energetically favorable to buckle and pay the bending energy cost of a plectoneme instead of increasing the twist energy of unbuckled DNA. Once the DNA buckles into plectonemes, an increase in linking number is accommodated by an increase in plectoneme size and plectoneme writhe that keeps the DNA twist unchanged and the restoring torque plateaus [24]. This behavior is shown by the *N* = 0 (blue) curve in Fig. 1 G.

As discussed above (Sec. I B), in the case of chromatin, positive twists injected into the relaxed state (with all negative nucleosomes) are accommodated by nucleosome chiral transitions. Consequently, we obtain a regime with zero restoring torque for positively twisted chromatin (Fig. 1 C, G). Consistent with the role of nucleosome chiral transitions in the emergence of this regime, the regime extends over larger ranges of positive supercoiling densities for higher nucleosome counts (Fig. 1 D, G and Fig. S7). The experimentally observed low torque valley near zero linking number is in accord with model predictions [18] (Fig. 1 G). For the case of negative excess linking number and for positive excess linking number beyond the buckling threshold (*i*.*e*., once the DNA has started to form plectonemes), the chromatin restoring torque response is similar to bare DNA (Fig. 1 G and Fig. S7). Note that the key feature of the chromatin torsional response— a regime with near zero restoring torque for a range of positive supercoiling density— is robust to variations in the amount of writhe accommodated by the different nucleosome chiral states (Fig. S4-S6).

Fig. 1 D provides an overview of the chromatin restoring torque as a function of the supercoiling density *σ* and nucleosome density *ψ*. Here, we use a higher extensible force (*f* ≈ 1.0 pN) which melts DNA at relatively lower negative supercoiling densities [24, 28]. The co-existence of melted and twisted DNA leads to a plateau in the negative torque (Fig. S7). A regime with a positive torque plateau is seen for positive supercoiling densities involving the co-existence of twisted and plectonemic DNA. While this regime is also seen both in bare DNA and in the case of chromatin, the onset of this regime in the case of chromatin occurs at higher values for *σ* for higher nucleosome densities. This effect is due to the ability of chiral nucleosome transitions to accommodate positive supercoiling thereby delaying DNA buckling. These two regimes of torque plateaus are in addition to the chiral-transition-driven regime of near-zero torque discussed earlier. The chromatin torsional response, as shown in Fig. 1 D, was used in the simulation of RNAP dynamics throughout this manuscript.

### D. Coupling chromatin torsional response with RNAP dynamics to model eukaryotic transcription

During transcription elongation, the RNAP must track the helical groove of the DNA, accumulating a rotational angle of *ω*_0_*x* when transcribing a DNA segment of length of *x* nm. Here, *ω*_0_ ≡ 2*π/h* ≈ 1.85 nm^−1^ is the linking number density in unstressed double-stranded DNA. If the genomic segment under transcription is torsionally constrained, this accumulated angle is partitioned between the rotation of the RNAP *θ* (and the associated nascent RNA) and the DNA twist at the site of the RNAP *ϕ*:

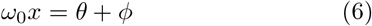

The angle *ϕ* determines the excess linking number injected into the genomic DNA and, thus, the restoring torque applied by the DNA or chromatin. Following the approach in [14], we write a torque balance equation for each RNAP:

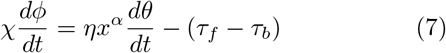

Here, *χ* is the DNA twist mobility, *η* is the coefficient of friction, and *α* is an exponent that determines how fast the viscous drag on the RNAP-nascent RNA complex grows with an increase in the nascent RNA length (which equals *x*, the distance moved by the RNAP). The term *ηx*^*α*^ thus determines the rotational mobility of the RNAP. *τ*_*f*_ and *τ*_*b*_ are the restoring torques applied by on the RNAP by the genomic segment downstream and upstream from the RNAP, respectively. While in the case of prokaryotes, *τ*_*f*_ and *τ*_*b*_ are only dependent on the excess linking number or supercoiling density in the respective genomic segments (*i*.*e*., *τ*_*f*_ ≡ *τ* (*σ*_*f*_) and *τ*_*b*_ ≡ *τ* (*σ*_*b*_)), in the case of eukaryotes, the restoring torque will also depend on the nucleosome density in the genomic segments, *i*.*e. τ*_*f*_ ≡ *τ* (*σ*_*f*_, *ψ*_*f*_) and *τ*_*b*_≡ *τ* (*σ*_*b*_, *ψ*_*b*_) (Fig. 1 D). The restoring torques applied by the chromatin segments on the RNAP were calculated using Eq. 5. Finally, the rate of RNAP translocation (*dx/dt*) is itself dependent on the net restoring torque acting on the RNAP with a torque-mediated stalling at *τ*_*c*_ = 12 pN *·* nm [14, 30]:

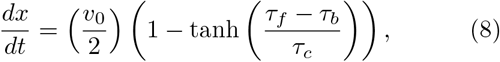

where the maximum RNAP velocity is *v*_0_ = 60 bp *·* s^-1^. Eq. 6-8 are solved to simulate the dynamics of a single RNAP.

To simulate transcription by multiple RNAPs, we consider a stochastic simulation setup wherein RNAPs are recruited to the transcription start site (TSS) at a rate *k*_*on*_ (Fig. 2 A). After recruitment, the dynamics of each RNAP is determined as described above. Supercoiling throughout the genomic segment is relaxed at a rate *k*_*relax*_, mimicking the activity of enzymes such as topoisomerases. Nucleosomes stochastically bind and unbind from the genomic segment at rates 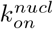 and 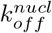, respectively, independent of the supercoiling density in the genomic segment.

**FIG. 2.**
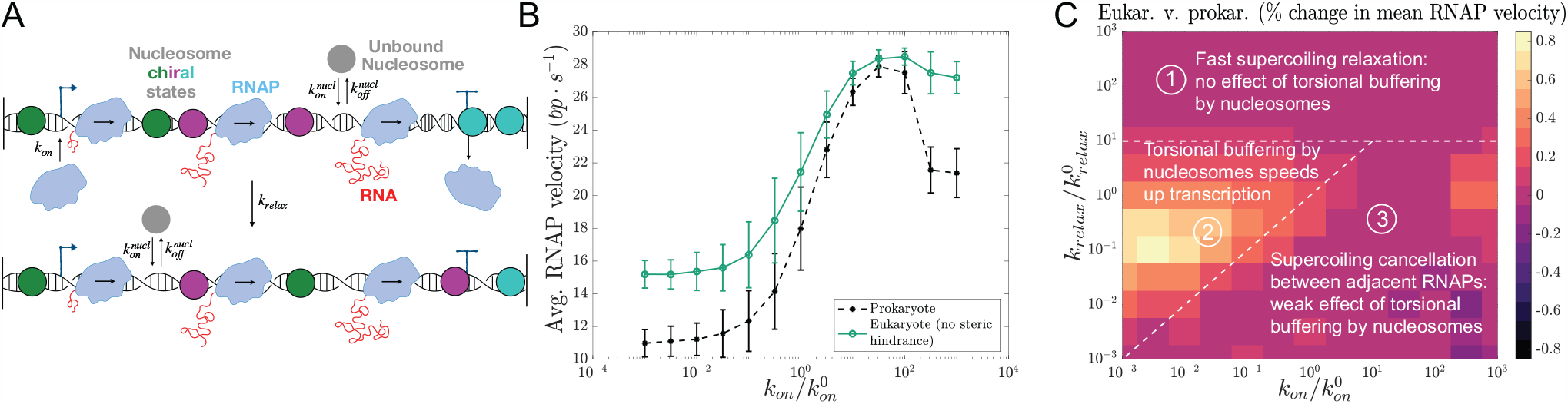
Effect of nucleosome-mediated torsional buffering on transcription elongation kinetics. **A** A schematic of the model for supercoiling-coupled transcription in the presence of nucleosomes (*i*.*e*., eukaryotic transcription). RNAPs are recruited to the transcription start site at a rate *k*_*on*_ while DNA supercoiling throughout the simulated genomic segment is relaxed at a rate *k*_*relax*_, mimicking DNA topoisomerase activity. Nucleosomes can bind to specific sites on the genomic segment at a rate 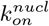 and unbind at a rate 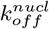. The movement of each RNAP is coupled to the restoring torques applied by the genomic segments upstream and downstream (Eqs. 6-8), building upon an approach previously utilized to analyze prokaryotic transcription with bare DNA torque response [14]. **B** The average RNAP velocity varies non-monotonically with *k*_*on*_ in both prokaryotes (without nucleosomes) and eukaryotes (with nucleosomes that do not sterically hinder RNAP movement). The presence of nucleosomes makes eukaryotic transcription elongation faster at low and high *k*_*on*_, while they are similar for intermediate values of *k*_*on*_. Error bars indicate the standard deviation. Nucleosome-driven weakening of chromatin torsional rigidity underlies the RNAP speed up in eukaryotes. **C** Percentage change in the average RNAP velocity in eukaryotes as compared to prokaryotes for different values of *k*_*on*_ and *k*_*relax*_. We indicate three distinct regimes. Regime 1: high topoisomerase activity where DNA torque-mediated constraints are minimal due to fast supercoiling relaxation, and hence torsional buffering by nucleosomes has no effect on transcription kinetics. Regime 2: torsional buffering by nucleosomes significantly speeds up eukaryotic transcription for genes with a lower initiation rate and lower topoisomerase activity. Regime 3: collective RNAP behavior, featuring supercoiling cancellation between adjacent RNAPs. Nucleosome-mediated torsional buffering has little effect in this regime since supercoiling-mediated RNAP slowdown is already being mitigated by their collective behavior. Here 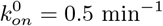 and 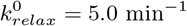 .

### E. Nucleosome-mediated torsional buffering speeds up transcription elongation

We used the above-described setup to simulate the transcription of a 5.3 kb gene (Fig. 2 and Fig. S8). We begin with the assumption that nucleosomes do not present any steric hindrance to RNAP movement, supported by previous studies reporting cooperative interactions between subunits of the RNAP complex and nucleosomes that can facilitate transcription through nucleosomes [31]. We find t hat t he a verage r ate o f transcription elongation, given by the RNAP velocity, varies non-monotonically with the rate of transcription initiation *k*_*on*_ (Fig. 2 B). The increase of RNAP velocity with an increased rate of initiation originates from the cancellation of supercoiling between adjacent RNAPs. This regime of collective RNAP behavior has been observed experimentally [3] and in our previous model of prokaryotic transcription [14], *i*.*e*., without nucleosomes. We find that the collective regime is not much perturbed by the presence of nucleosomes, rather, the low initiation regime is significantly affected (Fig. 2 B).

At low *k*_*on*_, on average, a single RNAP is transcribing the gene at any given time. In this regime, the transcription elongation rate in eukaryotes is higher since the presence of nucleosomes lowers the net restoring torque acting on the RNAP as compared to the prokaryotic case of bare DNA (Fig. 2 B). At higher *k*_*on*_, multiple RNAPs transcribe the gene simultaneously at any given time (see Fig. S10. We find that an RNAP transcribes faster if additional RNAPs are subsequently recruited to the TSS behind it (Fig. S9), which originates from supercoiling cancellation and lies at the crux of the RNAP cooperation. Supercoiling cancellation in this regime diminishes the effect of DNA or chromatin torsional response on transcription; consequently, the difference between prokaryotic and eukaryotic average RNAP velocities decreases, and the two approach one another (Fig. 2 B). Finally, at very high *k*_*on*_, we obtain a “traffic jam”-like regime, where the average RNAP velocity is likely to be determined by the translocation rate of the most downstream RNAP [14]. The translocation rate of this RNAP will depend on the restoring torque applied by the DNA or chromatin segment downstream from the gene body. The lower restoring torque in the case of chromatin underlies the higher average RNAP velocity of eukaryotes in this regime (Fig. 2 B).

In addition to *k*_*on*_, the average RNAP velocity is also dependent on the rate of supercoiling relaxation *k*_*relax*_, where faster relaxation speeds up transcription elongation [3, 14]. Comparing the average RNAP velocities in prokaryotes and eukaryotes in the *k*_*on*_-*k*_*relax*_ space, we identified three regimes with distinct behaviors (Fig. 2 C). At very high *k*_*relax*_ (regime 1), fast supercoiling relaxation makes DNA torque-dependent effects irrelevant and the RNAP velocities are the same in prokaryotic and eukaryotic cases. Similarly, in the regime of emergent collective behavior between co-transcribing RNAPs (regime 3), the RNAP velocities are similar for both prokaryotes and eukaryotes. Supercoiling cancellation between adjacent RNAPs makes the effects arising from the altered torsional response of chromatin less prominent. At low *k*_*on*_ and low *k*_*relax*_ (regime 2), the DNA or chromatin torsional response strongly influences RNAP translocation (Fig. 2 C). Consequently, the average RNAP velocity is higher in the eukaryotic case with nucleosomes buffering the restoring torque acting on the RNAPs.

### F. Steric hindrance from nucleosomes with slow turnover impedes transcription elongation

We now investigate the role of steric interactions between nucleosomes and RNAPs on transcription elongation. Fig. 3 shows the behavior when nucleosomes act as rigid barriers to RNAP movement and an RNAP must wait for the nucleosome downstream to unbind before it can move forward. As expected, the average RNAP velocity is very low at low rates of nucleosome unbinding from the genomic DNA. The average RNAP velocities at all *k*_*on*_ values increase for faster nucleosome unbinding, approaching the scenario with no steric hindrance at very fast nucleosome unbinding rates. Note that the emergent cooperation between co-transcribing RNAPs is present even with nucleosomes acting as steric barriers. This highlights that the cooperation regime is a key feature of transcriptional kinetics and is seen across contexts (Fig. 2 B and Fig. 3).

**FIG. 3.**
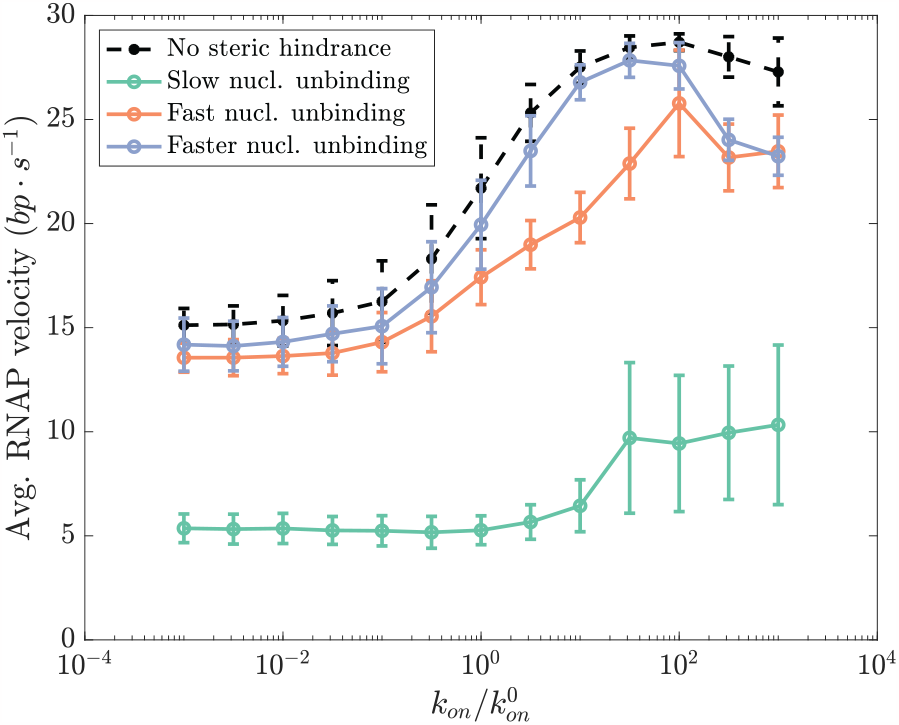
Effect of steric hindrance from nucleosomes on transcription elongation kinetics. Average RNAP velocity as a function of the transcription initiation rate (*k*_*on*_) for various nucleosome unbinding rates: slow 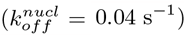, fast 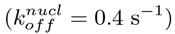, and faster 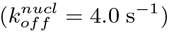. The nucleosome binding rate is kept unchanged in each case: 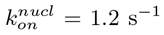. The dashed black line shows the case with no nucleosomal steric hindrance for comparison. Increased effective steric hindrance due to the slower unbinding of nucleosomes from the genomic DNA can decrease the average transcription elongation rate in eukaryotes. Since an RNAP must wait for the nucleosome in front of it to unbind before moving forward, the average RNAP velocity is lower at lower nucleosome unbinding rates. At higher nucleosome unbinding rates, the effective steric hindrance is lower, leading to kinetic behavior that resembles the no steric hindrance case.

Note that the treatment of nucleosomes as impenetrable barriers to RNAP movement is an extreme case. Experimental studies have shown that nucleosomes may need to only partially unbind from the DNA for the RNAP to pass through [31]. Thus, our results spanning the no steric hindrance to impenetrable barriers include the expected behavior *in vivo*.

### G. Predicting the transcription-dependent supercoiling profile in the yeast genome

We next simulated the transcription-supercoiling dynamics in long, multigenic segments of the budding yeast (*S. cerevisiae*) genome. The simulated segments were randomly chosen and ranged between 7 kb and 25 kb containing 4 to 25 genes. The *k*_*on*_ for each gene was chosen based on the gene expression level in the RNA-seq dataset from Guo *et al*. [5]. Fig. 4 shows the supercoiling density profiles over two multi-kilobase yeast genome segments as predicted by our model. The predicted super-coiling density profile is a function of the transcriptional state since the *k*_*on*_ rates for the various genes in the two segments are inputs to the model. Importantly, the density profile changes in response to perturbing the transcriptional state of one of the genes. Our model makes two testable predictions: first, the supercoiling density profile, and second, the change in the density profile upon perturbing the transcriptional (Fig. 4). Note that suppression (knockdown or KD) (Fig. 4 A) or overexpression (OE) (Fig. 4 B) of a gene can alter the supercoiling density profile not only in the neighborhood of the perturbed gene but also over large genomic neighborhoods.

**FIG. 4.**
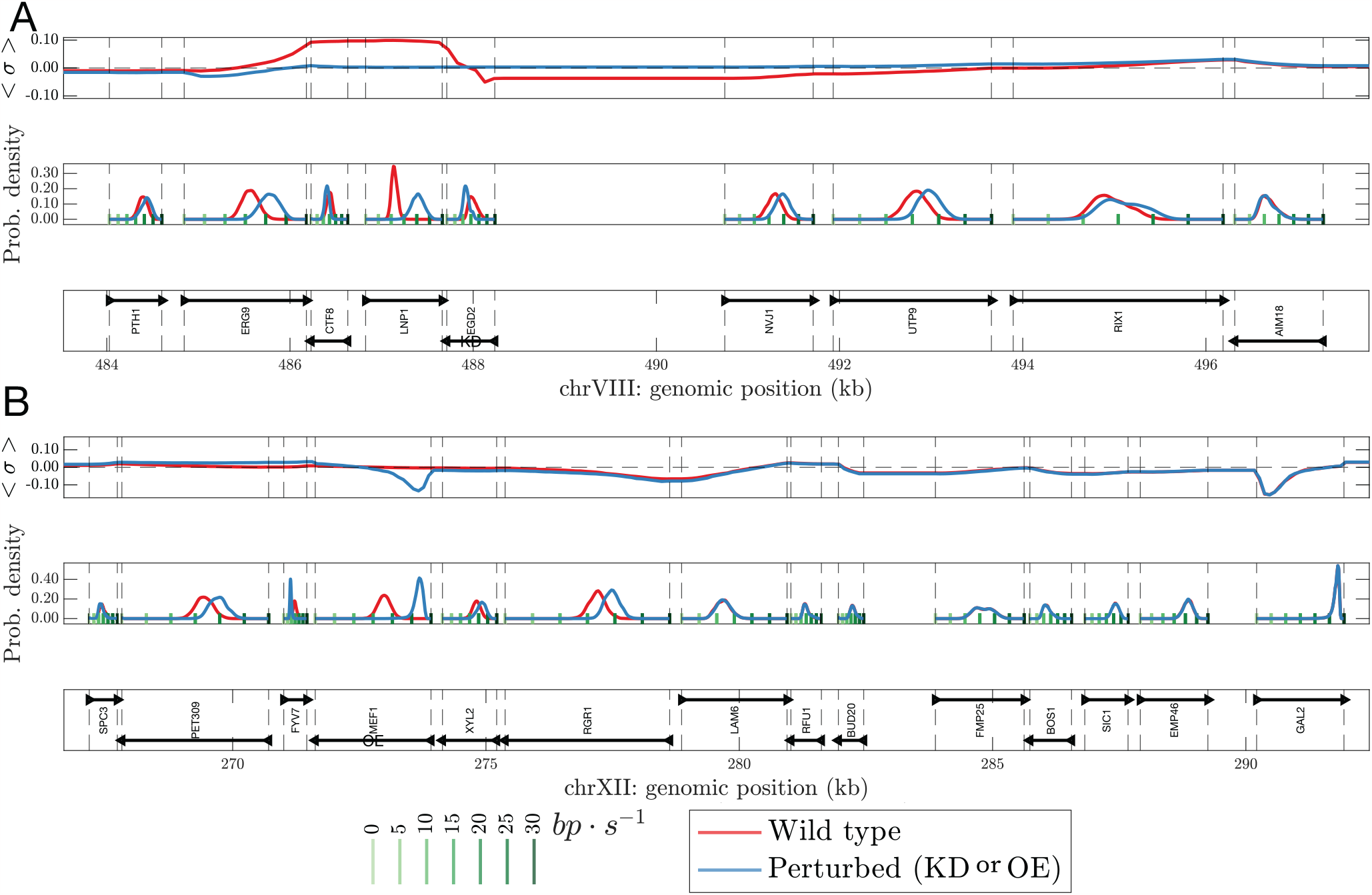
Transcription-generated supercoiling can perturb the elongation kinetics of neighboring genes. Two representative yeast (*S. cerevisiae*) genomic segments are shown in **A** and **B**. In each subplot, there are three panels. The top panel shows the supercoiling density, the middle panel shows the distributions of RNAP velocities for different genes, and the bottom panel shows the gene annotations for the segment. Using the RNA-seq data from Guo *et al*. [5], we set the *k*_*on*_ for each gene (Sec. SIV). We used our model to predict the “Wild-type” supercoiling density profile and RNAP velocities for the genes in each segment (shown in red). We additionally show a “Perturbed” phenotype for the supercoiling density profile and the RNAP velocities corresponding to a scenario where one of the genes in the segment is perturbed (knockdown of EGD2 in **A** and overexpression of MEF1 in **B**, shown in blue). Comparing the Wild-type with the Perturbed cases (red and blue curves), we see as expected, that perturbing a gene always has an effect on the local supercoiling density. Interestingly, however, the perturbation of the supercoiling profile may spread to longer distances, up to 10 kb or so, in a context-dependent manner. Note that while the example region in **A** shows a long-distance propagation of the supercoiling perturbation, the region in **B**, possibly due to its higher gene density stops the perturbation from spreading. The figure overall illustrates the capability of our framework to model transcription-supercoiling interplay for real genomic segments that are tens of kilobases long and contain multiple genes.

### H. Transcription-generated supercoiling as a mediator of inter-gene interactions

We probed the extent to which supercoiling-mediated interactions between neighboring genes can emerge in real genomic contexts. When a specific gene is perturbed, we find that the RNAP velocities of the neighboring genes are typically strongly affected (Fig. 4). For example, when EGD2 is knocked down, the average elongation rates of its immediate convergent (LNP1) and divergent (NVJ1) neighbors increase (note the shifts in the probability densities of RNAP velocities in Fig. 4). Interestingly, the effect of knocking down EGD2 is not only limited to its immediate neighbors: the average elongation rates for ERG9 and UTP9 (one gene away), and RIX1 (two genes away) change as well. However, not all genes one or two genes away are affected. This suggests that supercoiling-mediated effects may propagate through genes in a context-dependent manner. Our model can quantitatively predict supercoiling-dependent variations in the transcriptional kinetics of real gene clusters as well as synthetic constructs, such as a multi-gene plasmid [7, 32].

Experiments have shown, both in prokaryotes [3, 32] and eukaryotes [7], that transcription-generated supercoiling can affect the transcription kinetics of neighboring genes in a manner dependent on the relative orientation of the genes. We previously showed for prokaryotes that RNAPs transcribing neighboring genes oriented in tandem can cooperate, speeding up one another. In contrast, RNAPs co-transcribing genes in divergent and convergent orientations antagonize and slow one another down [14]. Since the qualitative coupling of RNAP translocation and DNA supercoiling in eukaryotes is the same as in prokaryotes, the qualitative rules for supercoiling-dependent neighbor interactions remain unchanged: activation for tandem and suppression for divergent and convergent orientations (Fig. 4 and Fig. S13). Note that we do not incorporate supercoiling-dependent variations in transcription initiation [15, 33], which is expected to make the gene interactions more nuanced and is left for future studies.

### I. Gene bodies show a gradient of supercoiling accumulation

Analyzing the genes within our simulated segments, we find that the variation of the supercoiling density along the gene body depends on the transcriptional state of the gene (Fig. 5). In the case of weakly expressed genes, there is minimal accumulation of DNA supercoils in the gene body. In the case of strongly expressed genes, negative supercoiling accumulates close to the transcription start site. Interestingly, the supercoiling density becomes more negative as one moves into the gene body, indicating the presence of highly untwisted DNA in the promoter-proximal part of the gene body. The supercoiling density then gradually becomes less negative towards the middle of the gene bodies and, eventually, positive close to the transcription end site. Since transcription over longer genomic distances generates more supercoiling, the gene body supercoiling density profile is further dependent on the gene length: longer genes accumulate more negative supercoiling close to the transcription start site as well as more positive supercoiling close to the gene end (Fig. 5).

**FIG. 5.**
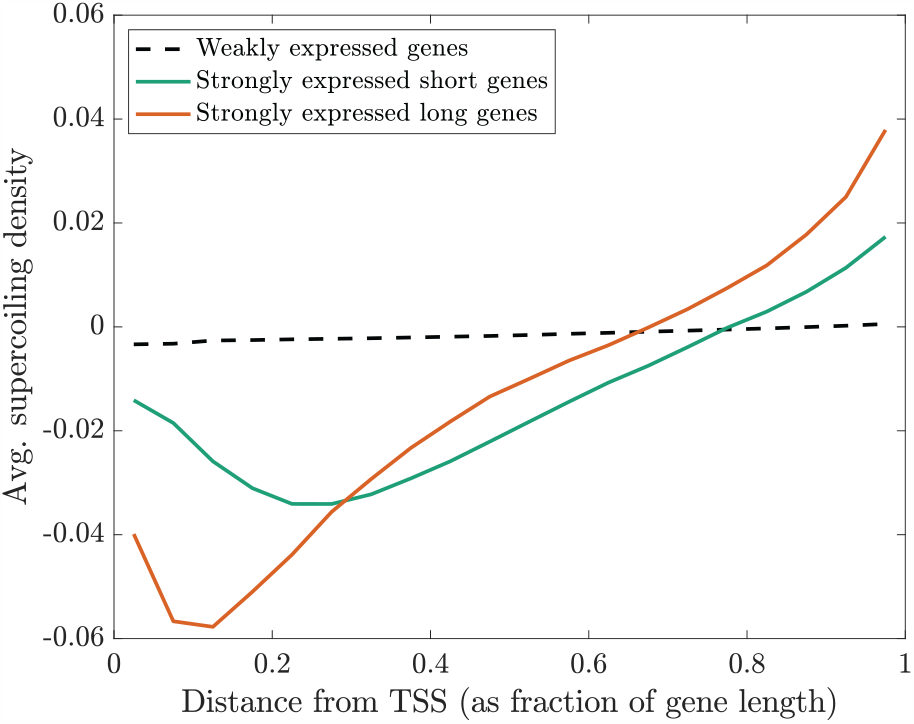
Supercoiling density profile in the bodies of yeast genes. Model prediction of the average supercoiling density in the gene body of yeast genes with different lengths and expression levels. The average was calculated over 68 weakly expressed genes 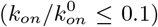, 22 strongly expressed genes 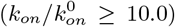 shorter than 0.5 kb, and 21 strongly expressed genes longer than 1.5 kb. Genes were assigned *k*_*on*_ values based on the RNA-seq data from Guo *et al*. [5] (Sec. SIV).

### J. Comparison with experiments

Guo *et al*. [5] developed GapR-seq, an assay for profiling the level of positive supercoiling genome-wide in both prokaryotes and eukaryotes. Applying this method to the budding yeast *Saccharomyces cerevisiae*, the study showed that the positive supercoiling accumulation was transcription-dependent. We simulated the supercoiling profile for 32 randomly chosen yeast genomic segments and compared it with the GapR-seq signal from Guo *et al*. [5] (Fig. 6). Note that while our model predicts the actual DNA supercoiling density, GapR-seq assay reports the relative abundance of positive supercoiling at a genomic locus.

**FIG. 6.**
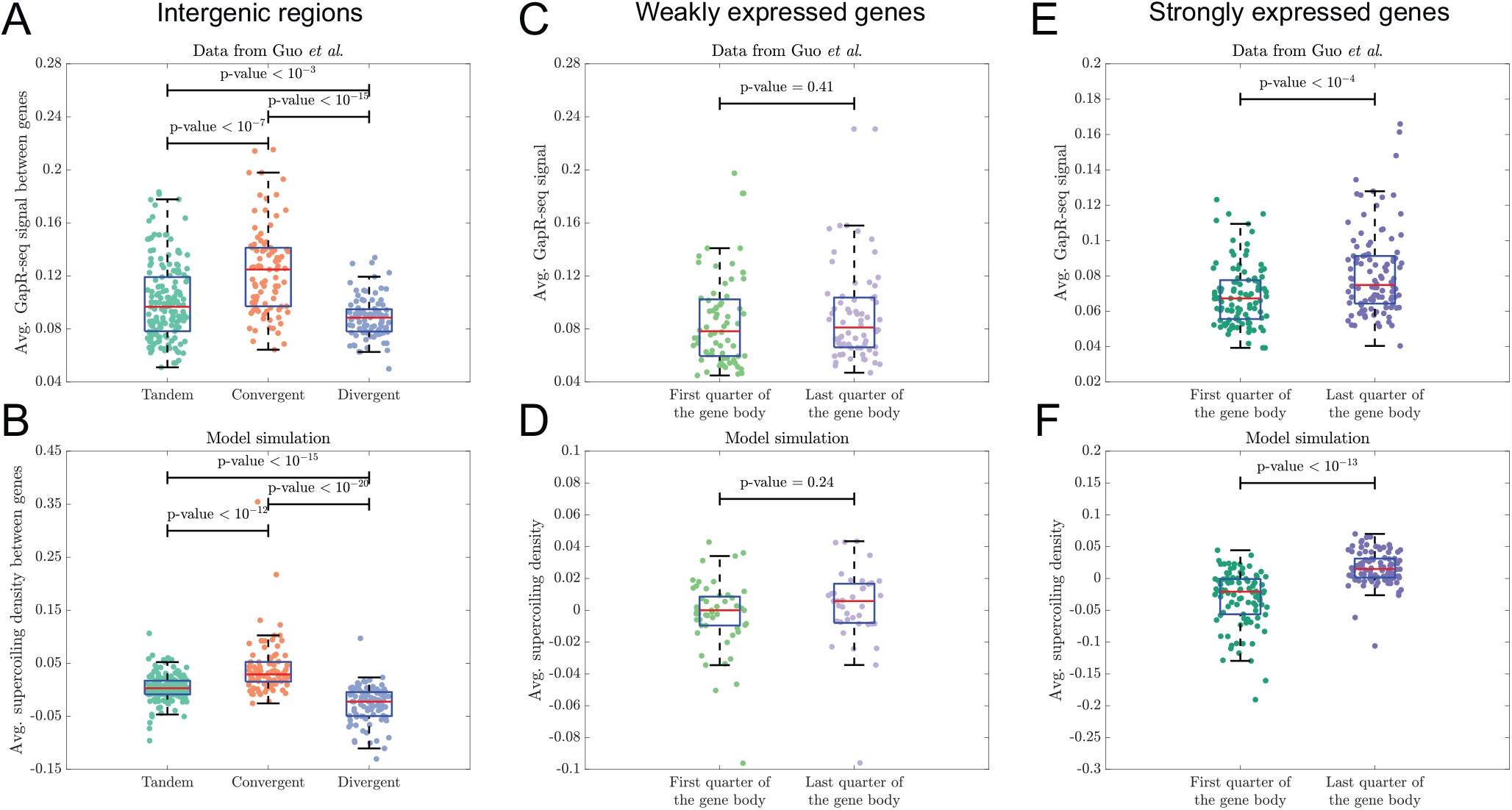
Comparison of supercoiling densities predicted by model simulations with GapR-seq data for yeast. **A** GapR-seq [5] data for intergenic regions shows a higher signal for intergenic regions between convergent genes as compared to regions between divergent or in tandem gene pairs, indicating higher accumulation of positive supercoils in the regions between convergent genes. A total of 351 intergenic regions are shown: 170 regions between genes in tandem, 89 regions between convergent genes, and 92 regions between divergent genes. **B** Model simulations for yeast genomic segments containing the genes in **A** recapitulate the trend in supercoiling densities shown in **A. C** In the case of weakly expressed genes 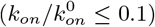, the GapR-seq signal shows no significant difference between the beginning and end of the gene bodies. 68 weakly expressed genes are shown here. **D** Model predictions of supercoiling densities recapitulated the trend shown in **C. E** In the case of strongly expressed genes 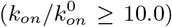, the GapR-seq signal indicated a higher accumulation of positive supercoils close to the end of the gene body. 102 strongly expressed genes are shown here. **F** Model predictions of supercoiling densities recapitulated the trend shown in **E**. The transcription initiation rates for the genes in our simulations were chosen based on the RNA-seq data from Guo *et al*. [5] in the same manner as for Fig. 4. These *k*_*on*_ values were used to classify the genes as weakly or strongly expressed. Yeast GapR-seq profile (data shown in panels A-C) was taken from the study by Guo *et al*. [5]. All p-values are for a two-sample *t* -test, with the null hypothesis that the data in the two groups are drawn from distributions with the same mean.

In agreement with the GapR-seq profiles, our simulations show that the extent of positive supercoiling is the highest in the intergenic regions between convergent genes and lowest in the regions between divergent genes (Fig. 6 A, B). Analysis of the GapR-seq signal in gene bodies showed that there is no significant difference in the average GapR-signal near the start and end of gene bodies for weakly expressed genes (Fig. 6 C). Whereas, for strongly expressed genes, positive supercoiling accumulated in the gene body close to the gene end (Fig. 6 E). Both these trends were recapitulated for the supercoiling profiles predicted by our model (Fig. 6 D, F). Overall, our model simulations recapitulate transcription-associated supercoiling features obtained from genome-wide positive supercoiling profiling in yeast.

## II. DISCUSSION

In the present study, we have developed a free energy minimization-based description of the chromatin torsional response (Fig. 1). Our model compares favorably with available experimental data and suggests chiral transitions by nucleosomes as the driver of the low torsional stiffness of the chromatin fiber (Fig. 1) [17, 18]. These chiral states, storing differing amounts of DNA writhe may interconvert via rotations about the dyad axis and accommodate DNA twists to weaken the torsional response(Fig. 1 and Fig. 7). We then integrated the chromatin torsion from this model into a previously proposed stochastic simulation framework [14] to investigate supercoiling-mediated aspects of transcription elongation kinetics in eukaryotes. Our major finding is that nucleosomes may have a dual effect on transcription. While binding to the gene body may hinder RNAP translocation, lowering the torsional stiffness of chromatin facilitates faster RNAP motion (Fig. 2 and Fig. 3). Using the model, we predicted the transcription-generated supercoiling profile in the yeast genome (Fig. 4). We find that genes may interact via DNA supercoiling, such that perturbation in the transcription state of a gene may significantly affect the RNAP motion for the genes in the neighborhood. Transcribed genes typically showed a negatively supercoiled transcription start site and a positively supercoiled transcription termination site (Fig. 5 and Fig. 7). We also found that the supercoiling accumulation in the intergenic regions depends on the relative orientation of the flanking genes (Fig. 4). Our results for supercoiling accumulation in the intergenic and genic regions are in agreement with the experimental observations (Fig. 6) [5].

**FIG. 7.**
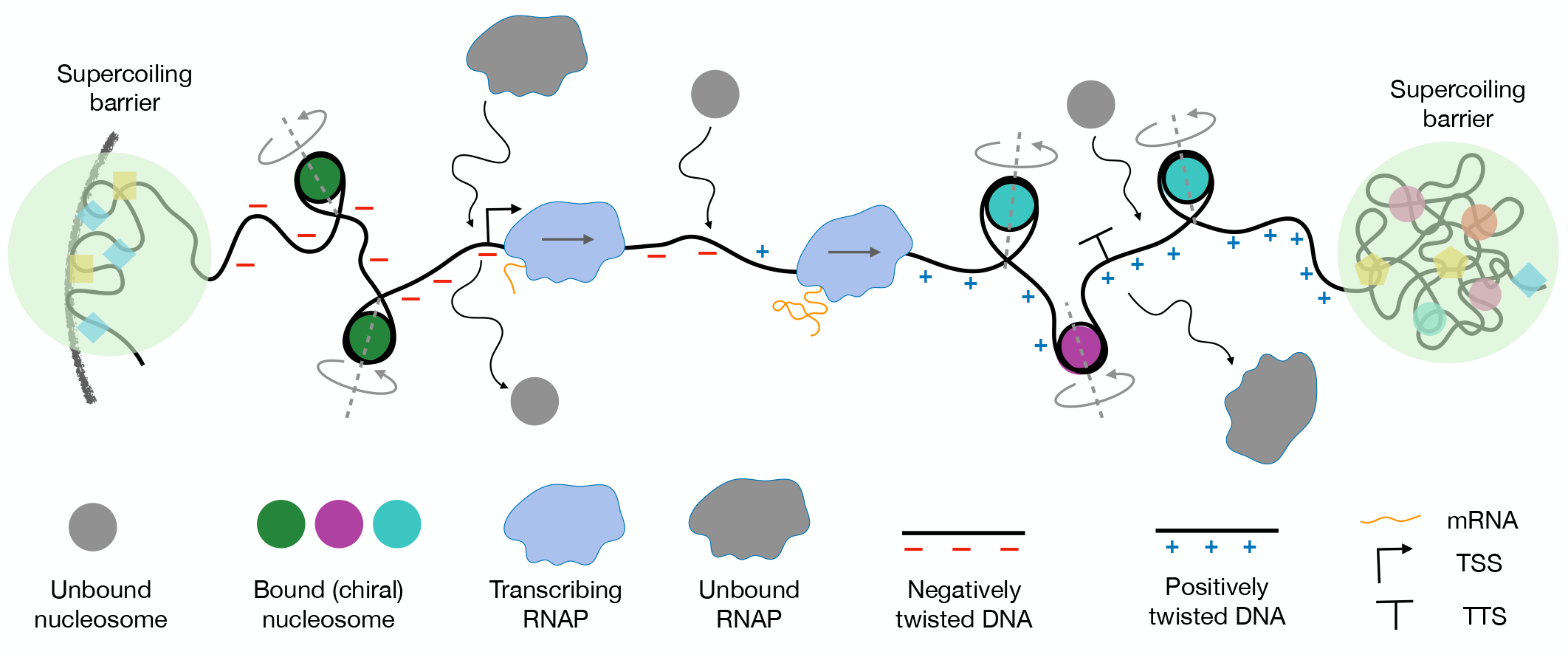
Graphical summary. Bound nucleosomes rotate to undergo chiral transitions and absorb DNA torque generated by RNAP translocation. The flanking region upstream (downstream) of a gene is typically negatively (positively) supercoiled, whereas the gene body shows a gradient of negative to positive supercoiling. Compact or cross-linked DNA globules or segments of DNA attached to the lamina could be possible barriers to supercoiling in vivo.

Our model simulations show that, just as in the prokaryotic case, co-transcribing RNAPs in eukaryotes can cooperate to speed up transcription elongation (Fig. 2). Such supercoiling-mediated cooperation, not requiring physical contact between adjacent RNAPs, has been experimentally confirmed in *Escherichia coli* [3]. We predict that such a cooperative regime would be prominent in eukaryotes (Fig. 2 B) as well, particularly under fast nucleosome turnover (Fig. 3). The gene orientation-dependent mechanical coupling of genes has also been observed both in prokaryotes and eukaryotes [3, 7]. Our model, incorporating the complex interplay between stochastic RNAP recruitment, supercoiling dynamics, and gene orientation, can serve as a useful framework for analyzing the complex behavior seen in experimental studies, and for identifying physiological regimes of interest.

The statistical mechanical model used to calculate the chromatin torsional response in the present study is simple. Contributions from nucleosome stacking [19, 34] or DNA sequence-dependence have been currently ignored. While the chiral transitions are central to the low torsional response, there may be a complex interplay between inter-nucleosome interactions and chiral transitions. Note that the kinetics of these chiral transitions may also be influenced by epigenetic modifications on histone tails [35]. Such considerations may be relevant to building more quantitatively accurate models as more experimental data become available.

Our model simulations can predict genomic supercoiling density profiles as a function of the transcriptional state (Fig. 4). The predicted supercoiling density profile may then be translated into predictions of nucleosomal conformations in different parts of the genome using our model of the chromatin torsional response (Fig. 1 C and Fig. S1. These predictions can be tested against nucleosome-level genomic structural features profiled by techniques such as Hi-CO [36] and RICC-seq [37]. We note that such predictions would benefit from a more detailed model of the chromatin free energy (see [38] for an example) such as one that incorporates higher-order chromatin structures [39, 40]. We assume these segments (typically 10-20kb long) to be insulated from a supercoiling perspective, which is in the same order of magnitude as bacterial supercoiled domains [41].

It has long been recognized that nucleosomes present a steric barrier to transcription, both *in vitro* [42] and *in vivo* [43, 44]. Our model of transcription elongation in eukaryotes shows that this inhibition is not the only mechanical effect of nucleosomes on transcription: nucleosomes can buffer RNAP-generated DNA torque and speed up transcription elongation. Thus, the overall effect of nucleosomes on the transcription elongation rate depends on the relative contribution from the two opposing effects (Fig. 3). Quantitative estimates concerning the nucleosomal barrier to RNAP movement are lacking. However, the fact that average transcription elongation rates in eukaryotes and prokaryotes are comparable would suggest that eukaryotic transcription operates in the regime of weak steric hindrance (or fast nucleosome unbinding; see Fig. 3). Multiple processes have been implicated in such modulation of the nucleosome barrier [22, 31]. The presence of the histone variant H2A.Z (instead of H2A) in nucleosomes has been shown to increase the nucleosome turnover rate, reducing the barrier to transcription [45, 46]. The histone chaperone FACT, which travels with the RNAP, can relieve RNAP stalling at nucleosomes by destabilizing histone-DNA contacts [47]. and promoting nucleosome eviction [48]. The various nucleosome remodelers, that use ATP to assemble, evict, or slide nucleosomes, also serve to alter the overall magnitude of the steric hindrance effect of nucleosomes on RNAPs [22]. These mechanisms of attenuating the nucleosome steric hindrance, along with RNAP speed-up from torsional buffering by nucleosomes, ensure fast transcription in eukaryotes. The modeling framework can be used to predict the qualitative effect of perturbing any of the aforementioned mechanisms.

Note that chromatin supercoiling can itself alter nucleosomal dynamics. Single-molecule assays have shown that nucleosome assembly is faster on negatively super-coiled DNA while positive supercoiling inhibits nucleosome binding [49]. A similar assay has shown that positive supercoiling can evict H2A / H2B dimers from nucleosomes, leaving behind tetramers [50]. Consistent with this observation, nucleosomes have been shown to be depleted from the region downstream of a highly transcribed gene in yeast [51]. In the present study, with a focus on the effect of torsional buffering on transcription elongation, we have simulated the simpler scenario where the nucleosome binding / unbinding kinetics are independent of the supercoiling density. Additionally, in contrast to previous theoretical studies [15, 33], we have assumed that transcription initiation (*i*.*e*. the model parameter *k*_*on*_) is not a function of the supercoiling density at the transcription start site. Both these dependencies may be incorporated into the approach described here and present promising future directions.

Comparing the predicted supercoiling density profiles in different genomic regions with the three-dimensional chromatin architecture of these regions obtained by Hi-C assays [52] is an exciting prospect. While it is not clear which elements constitute supercoiling barriers, threedimensional structures like compact globules or chromatin segments attached to nuclear bodies like lamina may act as barriers to twist diffusion since DNA may be heavily cross-linked in these regions. Diffusion of supercoils by rotation of these barriers is also a possibility that may be incorporated in the model. Although a connection between chromatin supercoiling and 3D chromatin architecture has been posited (for example, see Figure S2, panel J and the accompanying discussion in [53]), conclusive studies are lacking due to technical challenges like low resolution of supercoiling density genome-wide [54] or the inability to profile both positive and negative supercoiling levels [5]. Predicted transcription-dependent supercoiling profiles could help identify genomic regions where aspects of transcription, supercoiling, and 3D genome may be probed by targeted experiments [11]. The present model could further be extended to include additional biological processes that have been shown to exhibit supercoiling dependence such as the formation of R-loops [55] and recruitment of SMC complexes [56, 57]. Altogether, the model of supercoiling-transcription interplay described here can serve as a foundation for developing a DNA mechanics-based connection between genome architecture and cellular function.

## III.ACKNOWLEDGEMENT

This work was supported by the Center for Theoretical Biological Physics sponsored by the National Science Foundation (NSF) grant PHY-2019745, by NSF grants PHY-2210291 and DMR-2224030, and by the Welch Foundation (grant C-1792).

## SUPPLEMENTARY METHODS

Our modeling framework has two components: first, a statistical mechanical approach to recapitulate the torsional response of chromatin, and second, a framework to simulate transcription elongation wherein RNAP dynamics is coupled to the torsional mechanics of chromatin. The two components are described below.

### SI. Modeling the chromatin torsional response

We model the torsional response of chromatin by including the energetic and topological contributions from nucleosome binding [17, 18] in a twistable worm-like chain model of naked DNA [29]. As mentioned previously in Sec. I A, we consider a chromatin segment with DNA of length *L*_0_ and *N* nucleosomes, under an extension force *f* (Fig. 1 A). Each nucleosome is treated as a structural unit that absorbs *ℓ* = 60 nm (or 177 base pairs) of DNA. Nucleosome binding to DNA is stabilized by a negative binding energy *ϵ* = − 30 *k*_*B*_*T* . Since DNA wraps around each nucleosome in a left-handed fashion, nucleosome binding imparts an overall negative linking number in the form of writhe (Wr) to the chromatin segment [17–19]. Nucleosomes are not rigid objects and can exhibit structural [17] changes in response to extrinsic forces and torques. Here, we consider three structural states of nucleosomes (shown in Fig. 1 B) that differ in the relative orientation of the two linker DNA segments exiting the nucleosome-DNA complex, and thus the amount of linking number stored as writhe. The DNA not absorbed by wrapping around nucleosomes may be partitioned between a force-stretched state (of length *L*_*s*_) and a plectonemically buckled state (of length *L*_*p*_): *L*_0_ = *L*_*s*_+*L*_*p*_+*Nℓ*. When the chromatin is twisted, either by magnetic or optical tweezers in single-molecule studies [58], or by molecular motors such as RNA polymerases *in vivo*, the partitioning of the excess DNA linking number ∆Lk is governed by Eq. 3.

The total free energy of chromatin, as shown in Eq. 2, is computed by summing the contributions from the extended state (*F*_*s*_), the plectoneme state (*F*_*p*_), and the nucleosome states (*F*_nuc_). The extended state energy *F*_*s*_ comes from the force-extension energy that has non-linear entropic elasticity and the quadratic DNA twisting energy [24, 29]:

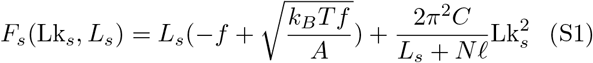

Here, *A* = 50 nm is the DNA bend persistence length and we assume a bare DNA twist stiffness of *C* = 100 nm [29] for DNA twisting within the nucleosome states. The plectoneme state energy *F*_*p*_ is given by a harmonic dependence on the linking number, similar to the extended state:

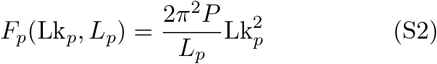

The twist modulus in the plectoneme state, *P* = 25 nm [24], is much lower than that for bare DNA (*C*), due to the screening of twist by plectoneme writhe. Finally, the nucleosome state energy is given by *F*_nuc_ = *Nϵ*, where *ϵ* = −30*k*_*B*_*T* is the DNA binding energy per nucleosome. Using different binding energies for the different nucleosome chiral states while keeping the differences between the states small does not significantly alter the chromatin torsional response (Fig. S1-S3).

We next construct a partition function incorporating all possible plectoneme length and nucleosome state configurations consistent with the linking number constraint given by Eq. 3. The chromatin torque *τ* can then be calculated using Eq. 5 while the end-to-end extension *z* for a given ∆Lk (as plotted in Fig. 1) can then be obtained from the partition function. Other averaged quantities may also be calculated from the partition function using standard procedures [25], as follows:

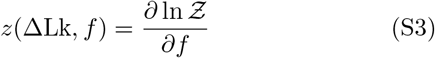

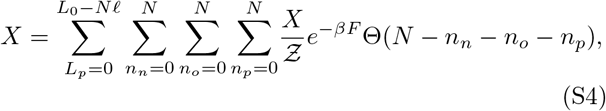

Here *X* denotes observables like the average number of open, positive, or negative nucleosomes (Fig. S1).

In Eq. 4, we assume that the total number of chromosomes is fixed; nucleosomes can only undergo chiral transitions to minimize free energy in response to torsional stress. Thus, Eq. 5 gives the torque as a function of the supercoiling density and the nucleosome count (or nucleosome density *ψ*), *i*.*e. τ*≡ *τ* (*σ, ψ*). This chromatin torsional response, shown in Fig. 1 D, is used in the model of RNAP dynamics described below.

#### Finite-size correction to torque calculation

Note that Fig. 1 D shows the torsional response in the thermodynamic limit, *i*.*e*. for a long genomic segment. Given DNA’s bending stiffness, a shorter DNA segment is likely to form plectonemes only at higher supercoiling densities. We incorporated this effect by including, in our model, a phenomenological dependence of onset of plectoneme formation on the length of the DNA segment. Let 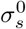 be the supercoiling density beyond which a long DNA segment starts forming plectonemes. Then, a DNA segment of length *l* will form plectonemes for *σ > σ*_*s*_(*l*) where

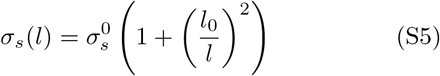

Here, *l*_0_ = 340 nm (or 1000 bp).

### SII. A model of DNA supercoiling-coupled transcription in eukaryotes

We adapt the model of transcription-supercoiling interplay in prokaryotes described previously by Tripathi *et al*. [14] to the case of eukaryotic transcription by incorporating the chromatin torsional response calculated in the previous section. From Eq. 6 and Eq. 7, we can write:

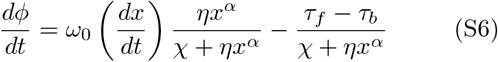

Eq. S6 and Eq. 8 can then be solved numerically to simulate the dynamics of a single RNAP. We used the following parameters for the simulations in this manuscript (same as in [14]): *χ* = 0.05 pN nm s, *η* = 5.0*×* 10^−4^ pN nm^-2^ s, and *α* = 1.5. The choice of these parameters was within the biophysical range; a detailed description of the rationale behind parameter choice can be found in [14].

In the case of eukaryotes, the restoring torques *τ*_*f*_ and *τ*_*b*_ are functions of the supercoiling density *σ* and the nucleosome density *ψ* in the corresponding genomic segment. We consider a genomic segment of length *L* extending from *X* = 0 to *X* = *L* with *M* RNAPs present at *X*_1_, *X*_2_, …, *X*_*M*_ . Let *ϕ*_*i*_ and *ϕ*_*i*+1_ be the DNA rotation angles at *X*_*i*_ and *X*_*i*+1_, respectively. Then, the supercoiling density in the segment bounded by the *i*^*th*^ and (*i* + 1)^*th*^ RNAPs is calculated as:

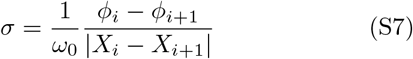

Here, we assume that the supercoiling density in a genomic segment depends only on the DNA rotation angle at the two ends of the segment, *i*.*e*. any twist generated at the ends of the segment diffuses instantaneously throughout the length of the segment (or, diffuses at times scales much faster than those associated with RNAP dynamics). To model transcription in torsionally constrained genomic segments, we choose the boundary conditions *ϕ*(*X* = 0) = *ϕ*(*X* = *L*) = 0.

Note that the nucleosome density *ψ* is defined as the fraction of DNA in the segment that is wrapped around nucleosomes, *i*.*e. ψ* = *Nℓ/L*_0_ where *N* is the number of nucleosomes in the segment at any given instant and *ℓ* = 60 nm.

The *σ* and *ψ* calculated for each genomic segment were used as inputs for the *τ* calculation scheme described above, and these *τ* values were used in Eq. 7 and Eq. 8. Note that simulating RNAP dynamics requires torque calculation every time the position of one or more RNAPs is updated. Therefore, carrying out the free energy minimization procedure to calculate *τ* each time would be prohibitively slow. To speed up the simulations, we used a 2D linear interpolation function fitted to the *τ* (*σ, ψ*) function shown in Fig. 1 D. The interpolation procedure was implemented using the C++ library linterp [59].

### SIII. Simulating eukaryotic transcription

Following the approach in Tripathi *et al*. [14], we simulate transcription by multiple RNAPs by setting up a stochastic simulation framework wherein RNAPs are recruited to the transcription start site at a rate *k*_*on*_ and supercoiling in the genomic segment is relaxed globally at a rate *k*_*relax*_: *ϕ*_1_ = *ϕ*_2_ = … = *ϕ*_*M*_ = 0 for RNAPs 1 … *M* . This setup is adapted for the eukaryotic case by including two additional events to model nucleosome turnover: binding and unbinding of nucleosomes from fixed DNA sites at rates 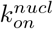and 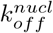, respectively (Fig. 2 A). Unless mentioned otherwise, we used 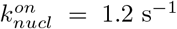 and 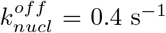 for all simulations. We incorporate the steric effect of nucleosomes on transcription by treating nucleosomes as rigid barriers to RNAP movement, *i*.*e*., if an RNAP encounters a nucleosome in its path, it must wait for the nucleosome to unbind before translocating further along the DNA. Additionally, RNAPs act as steric barriers to nucleosome binding: a nucleosome cannot bind to a DNA site that is occupied by an RNAP. Note that, in our setup, nucleosomes do not act as barriers to transcription initiation: if a nucleosome is present at the TSS when an RNAP recruitment event occurs, the nucleosome is dislodged before the RNAP binds the TSS. Throughout this manuscript, we make the simplifying assumption that the nucleosome dynamics are independent of the supercoiling density, *i*.*e*., 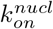 and 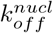 are not functions of *σ*. RNAP recruitment to the transcription start site is also assumed to be independent of the super-coiling density at the corresponding genomic locus.

The model dynamics were simulated using the Gillespie algorithm [60]. For every simulation setup in this manuscript, 16 independent runs were carried out. The RNAP velocities reported in the various figures were aggregated across the independent runs. To obtain the average number of co-transcribing RNAPs shown in Fig. S10 and the average nucleosome occupancies shown in Fig. S11, we sampled the system state at randomly chosen time points for each independent run. The averages shown were calculated from the sampled points across the 16 independent runs. To obtain the supercoiling density profiles shown in Fig. 3 and Fig. 6, we probed the supercoiling density at intervals of 34 nm (or 100 bp) on the genomic segment. The average supercoiling densities shown were then obtained by sampling the density at randomly chosen time points across the 16 independent runs. In Fig. 6 D, F, the average densities shown for the first and last quarters of the gene body were obtained by averaging over the points probed in each quarter. In Fig. 5, we probed the supercoiling density at 50 equally-spaced points in each gene body, independent of the gene length. Thus, the probed points were more closely spaced in the case of shorter genes. Once again, the average supercoiling densities shown were obtained by sampling the density at randomly chosen time points across the 16 independent runs.

### SIV. Simulating transcription of yeast genomic segments

We simulated transcription of 32 randomly chosen genomic segments of the budding yeast *Saccharomyces cerevisiae*. Results of these simulations are shown in Fig. 4-6. For these simulations, the *k*_*on*_ values for the various genes were chosen based on the RNA-seq data available from the study by Guo *et al*. [5] (Gene Expression Omnibus (GEO) [61] accession ID GSM5001912). For each gene, the overall expression level was calculated by averaging the RNA-seq signal in the gene body (non-normalized number of reads available from GEO); the expression level was then log-transformed, defining an RNA-seq signal, *sig*_RNA-seq_, for each gene. Thereafter, *k*_*on*_ of each gene was chosen as follows:

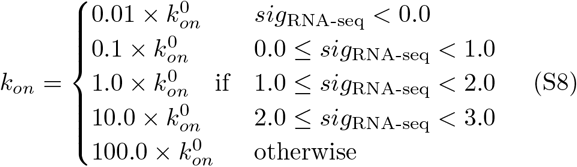

where 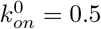.

A list of the yeast genomic segments simulated is available from https://github.com/st35/nucleosomes-supercoiling/tree/main/yeast_setup/configs. In each file available at this URL, first column is the gene name, third column is the gene length, fourth column is the gene strand (+1 for positive strand, −1 for negative strand) and the last column is 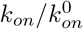 for the gene. These .config files can be directly used as inputs by our simulation code. The GapR-seq data shown in Fig. 6 was downloaded from GEO (accession ID GSM5001899).

### SV. Code availability

The python code used for free energy minimization to calculate the torsional response of DNA in the presence of nucleosomes is available online at https://github.com/s-brahmachari/Chromatin-supercoiling-and-transcription.

The C++ code used to simulate the transcription-supercoiling interplay is available online at https://github.com/st35/nucleosomes-supercoiling.

## SUPPLEMENTARY FIGURES

**FIG. S1.**
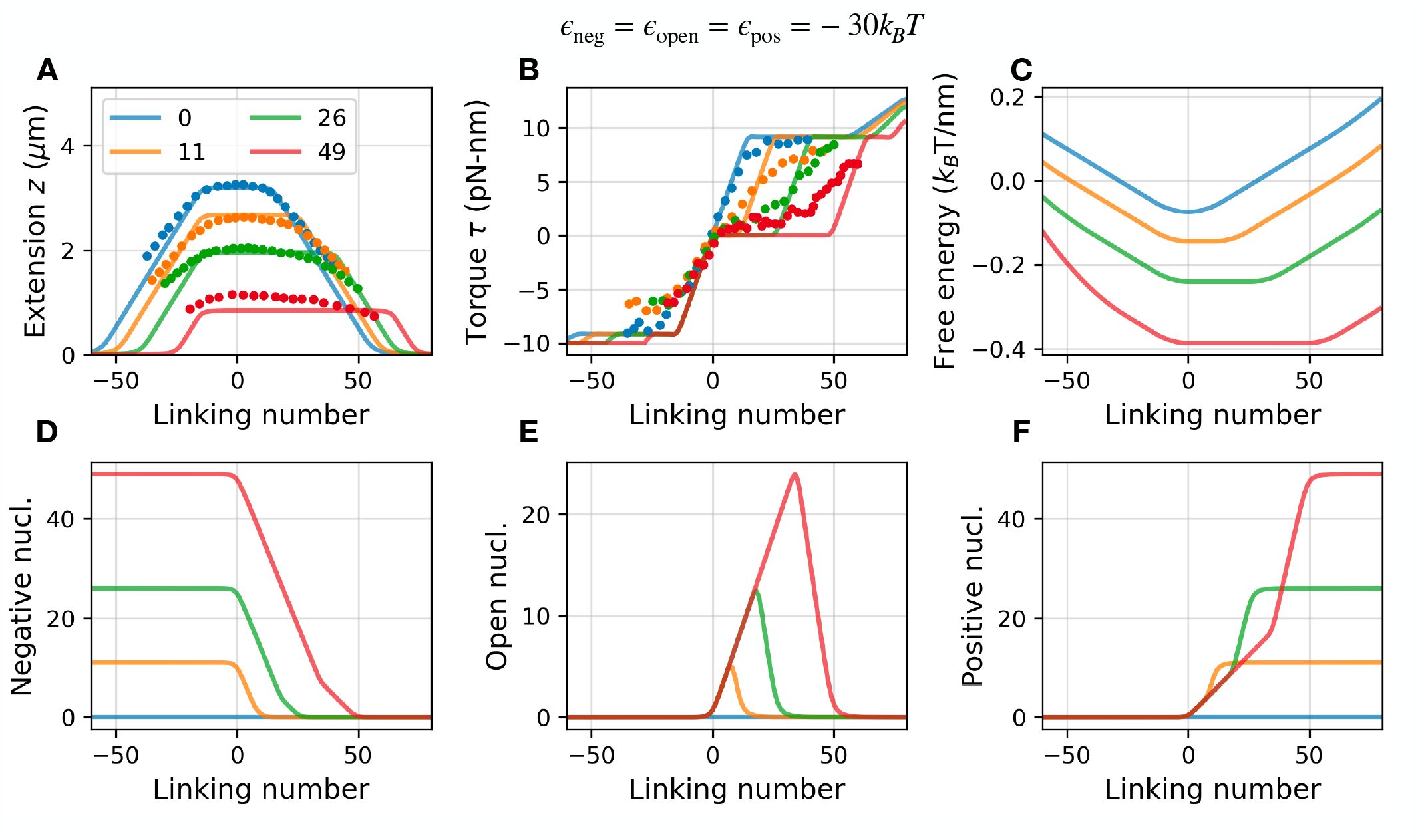
Torsional response of chromatin fiber with fixed binding energies for chiral states. (A) Extension, (B) Torque, (C) Free energy, (D) Number of negative nucleosomes, (E) Number of open nucleosomes, and (F) Number of positive nucleosomes as a function of excess linking number being injected into the chromatin fiber. We used a reference of Wr_ref_ = *N* Wr_*n*_ and the dots are experimental data from Le *et al*. [18]. The binding energy of all the nucleosomes are assumed to be fixed at =-30 *k*_*B*_ *T* . Note that the open nucleosome states are only transiently populated before switching to either positive or negative nucleosomes.

**FIG. S2.**
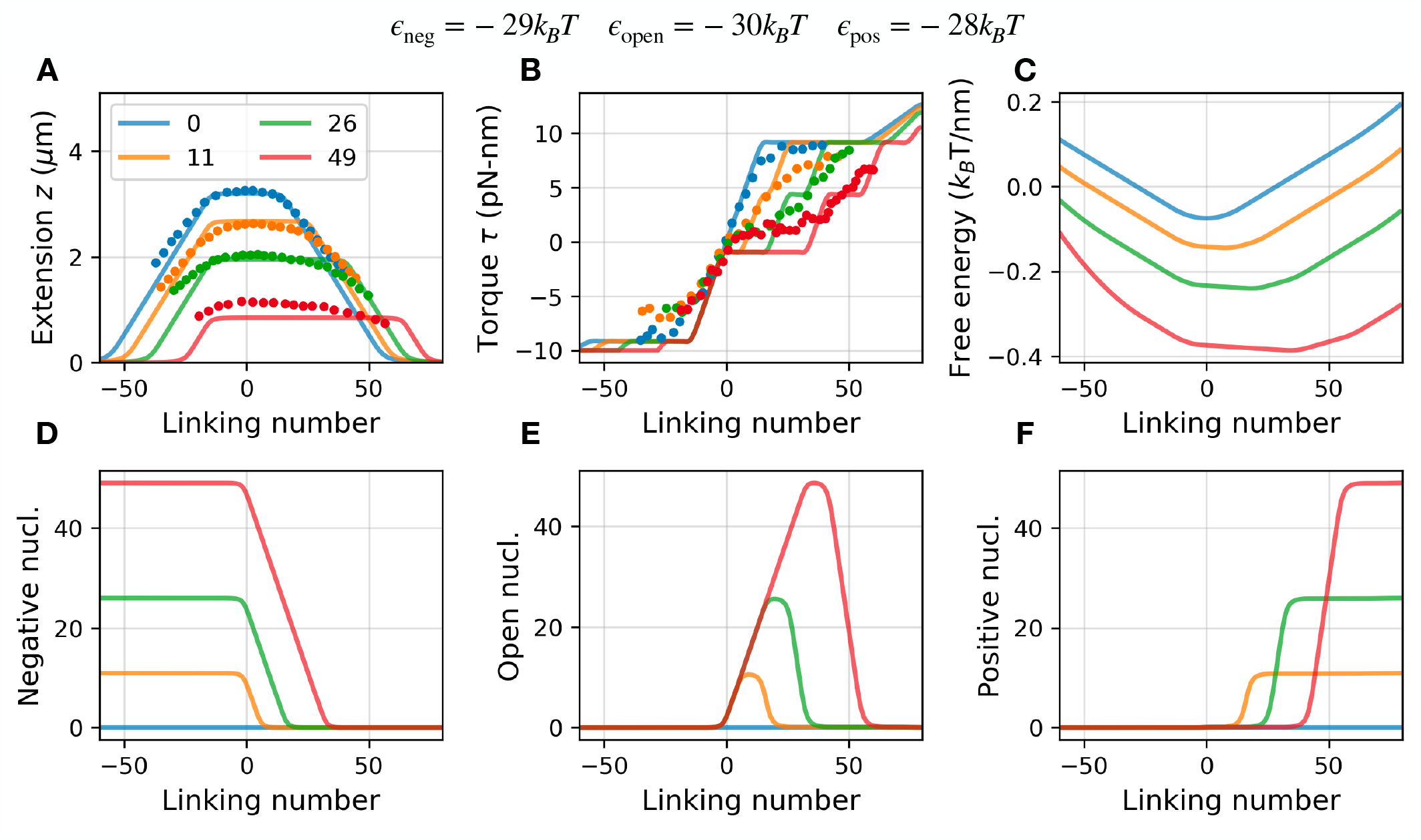
Torsional response of chromatin fiber with different (high) binding energies for chiral states. (A) Extension, (B) Torque, (C) Free energy, (D) Number of negative nucleosomes, (E) Number of open nucleosomes, and (F) Number of positive nucleosomes as a function of excess linking number being injected into the chromatin fiber. We used a reference of Wr_ref_ = *N* Wr_*n*_ and the dots are experimental data from Le *et al*. [18]. The binding energy of open states are favored compared negative and positive nucleosomes. We used *ϵ*_*open*_ = −30*k*_*B*_ *T, ϵ*_*negative*_ = −29*k*_*B*_ *T*, and *ϵ*_*positive*_ = −28*k*_*B*_ *T*, in accord with [17, 27]. Note that the open nucleosome states are stable and this introduces an additional kink in the torque plot. However, we argue that the overall torque response is similar for the purpose of transcription simulations.

**FIG. S3.**
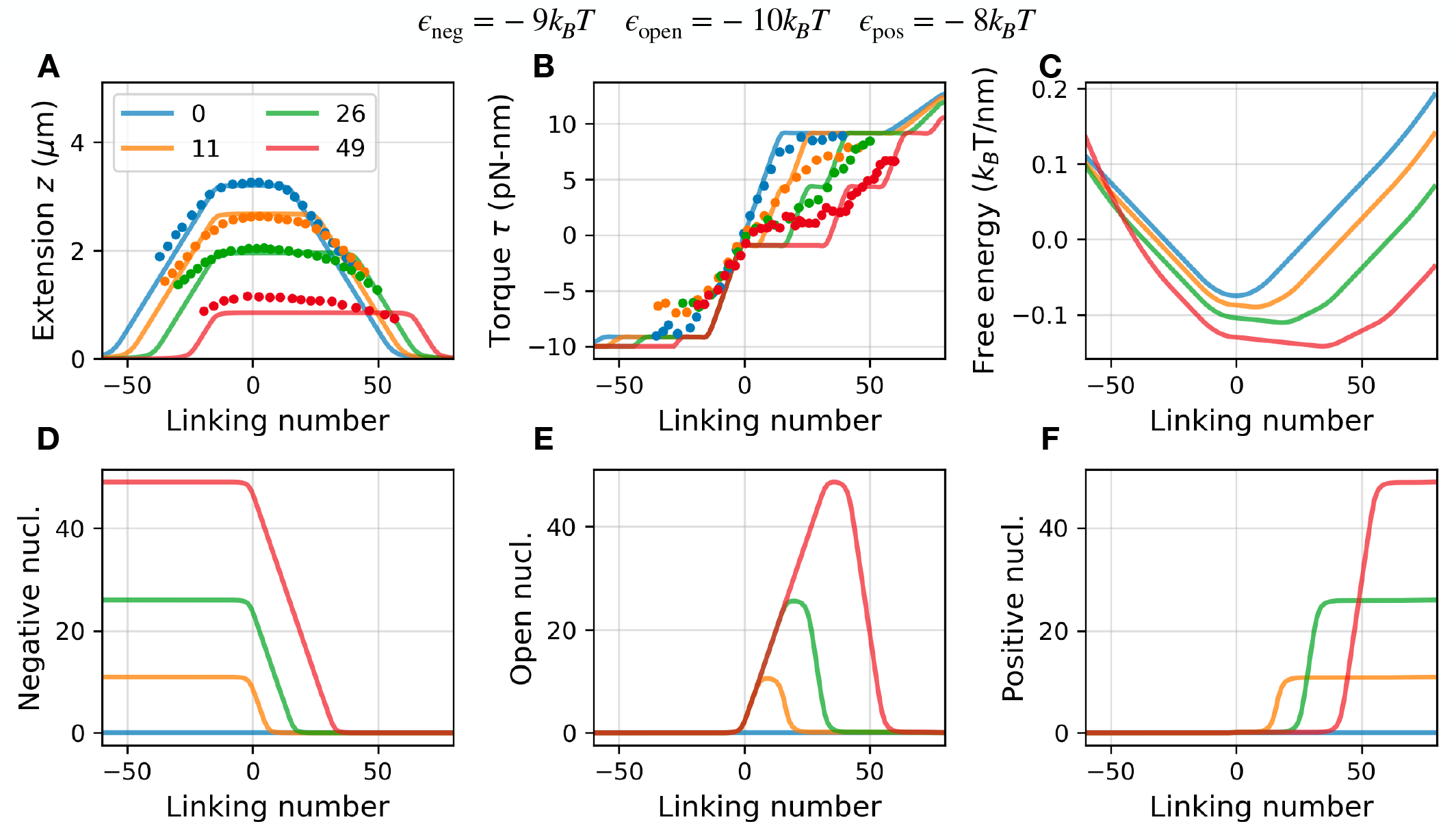
Torsional response of chromatin fiber with different (low) binding energies for chiral states. (A) Extension, (B) Torque, (C) Free energy, (D) Number of negative nucleosomes, (E) Number of open nucleosomes, and (F) Number of positive nucleosomes as a function of excess linking number being injected into the chromatin fiber. We used a reference of Wr_ref_ = *N* Wr_*n*_ and the dots are experimental data from Le *et al*. [18]. We keep the relative difference in the binding energies in accord with previous works [17, 27], but changed the overall binding energies, such that, *ϵ*_*open*_ = −10*k*_*B*_ *T, ϵ*_*negative*_ = −9*k*_*B*_ *T*, and *ϵ*_*positive*_ = −8*k*_*B*_ *T* . However, since our calculations are at a fixed nucleosome number, this leads to a constant shift in the free energy and has no effect on the torsional response.

**FIG. S4.**
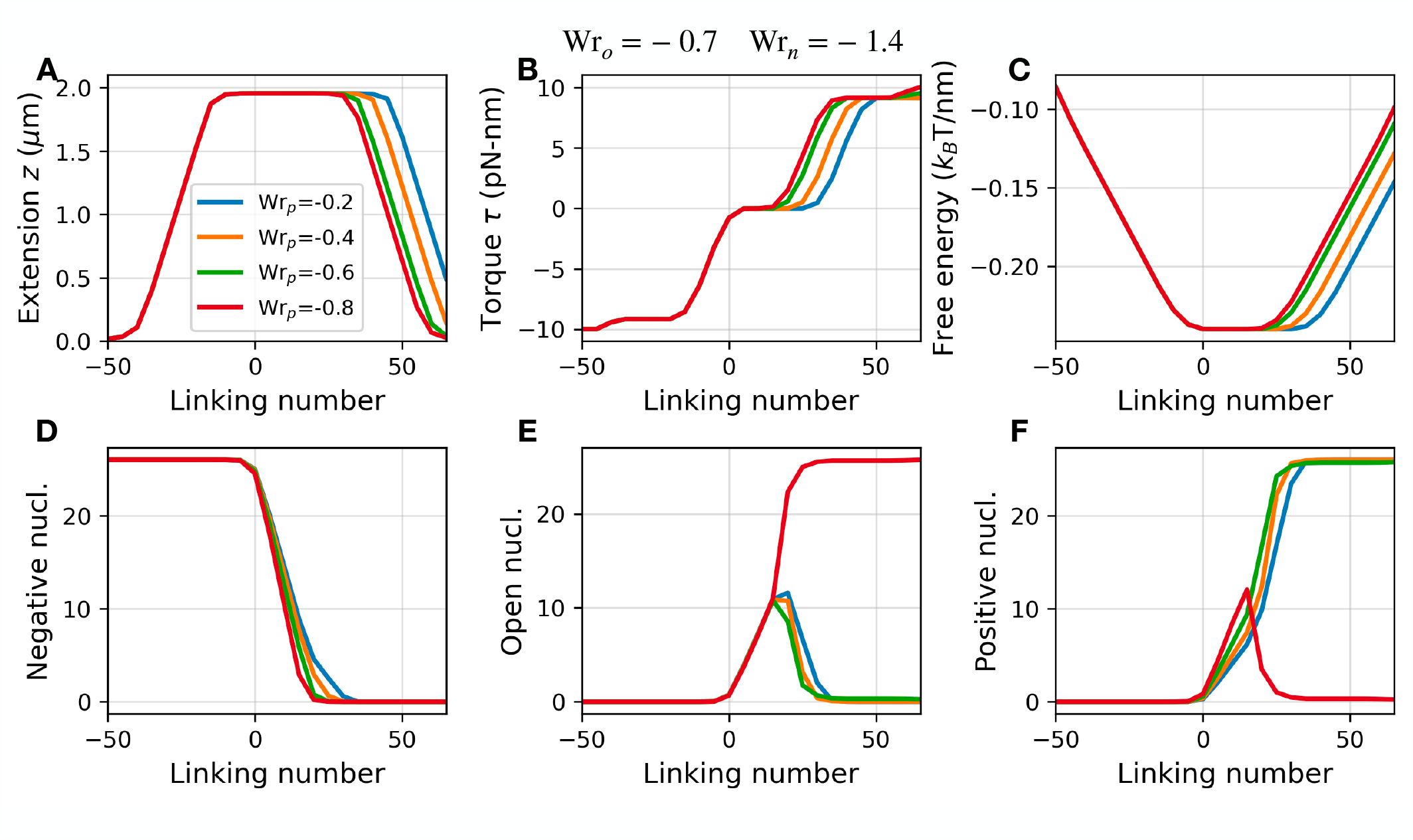
Torsional response of chromatin fiber with different writhe for positive nucleosome states. (A) Extension, (B) Torque, (C) Free energy, (D) Number of negative nucleosomes, (E) Number of open nucleosomes, and (F) Number of positive nucleosomes as a function of excess linking number being injected into the chromatin fiber. We used a reference of Wr_ref_ = *N* Wr_*n*_, corresponding to Fig. S1. Changing the writhe to less negative values increases the torque plateau because chiral transitions can absorb larger linking numbers. Note that the positive state is destabilized by the open state when the writhe of the positive state is more negative than the open state (Wr_*p*_ = −0.8). This, however, is a geometrically unphysical state if the nucleosome’s inner turn is not perturbed.

**FIG. S5.**
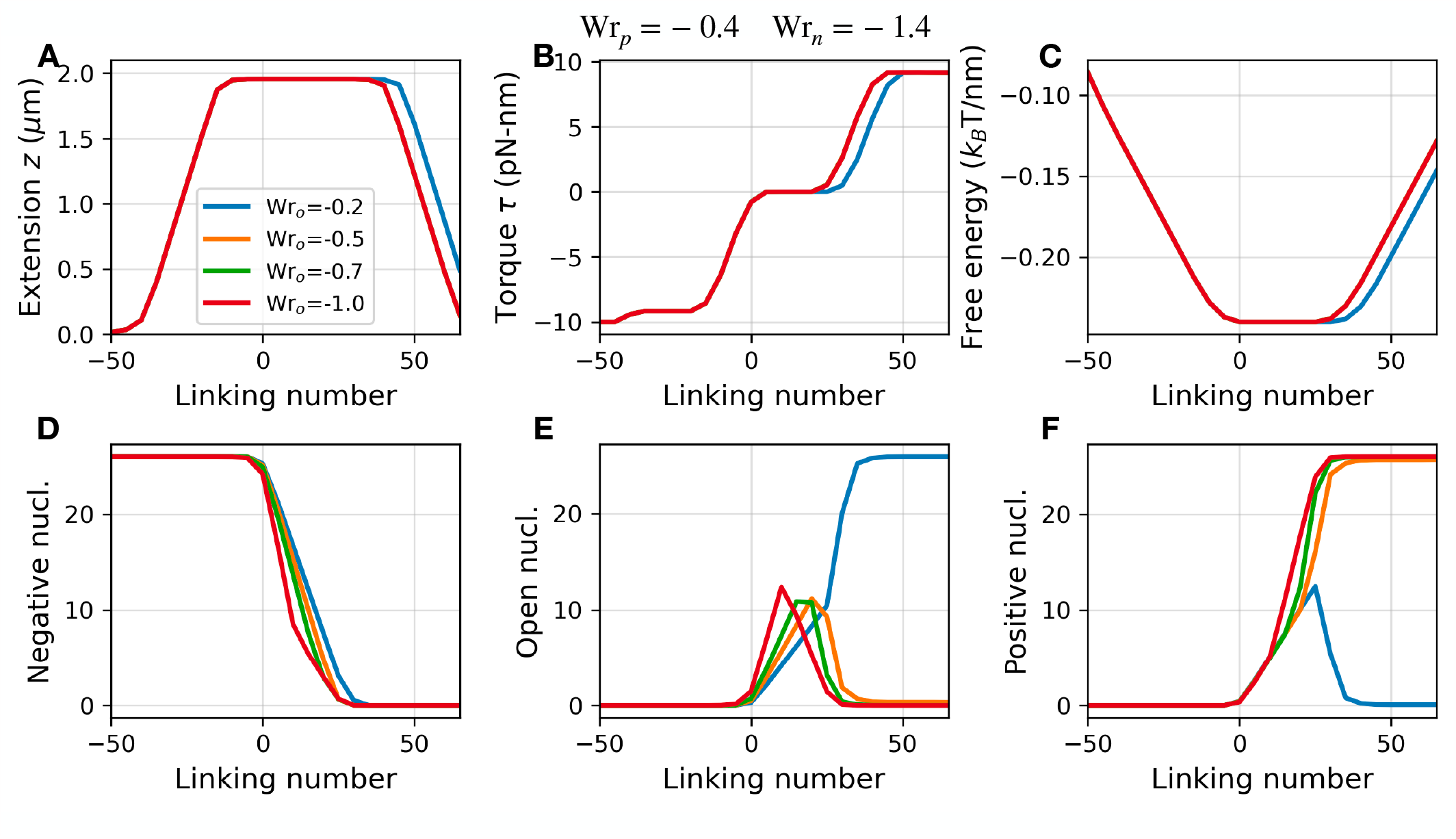
Torsional response of chromatin fiber with different writhe for open nucleosome states. (A) Extension, (B) Torque, (C) Free energy, (D) Number of negative nucleosomes, (E) Number of open nucleosomes, and (F) Number of positive nucleosomes as a function of excess linking number being injected into the chromatin fiber. We used a reference of Wr_ref_ = *N* Wr_*n*_, corresponding to Fig. S1. Changing the writhe has no effect on the torque or extension until the writhe exceeds the bounding positive state. When the open state stores less negative writhe than the positive state (Wr_*o*_ = −0.2), the positive state is destabilized and the open state dominates in the positive supercoiling regime. This, however, is a geometrically unphysical state if the nucleosome’s inner turn is not perturbed.

**FIG. S6.**
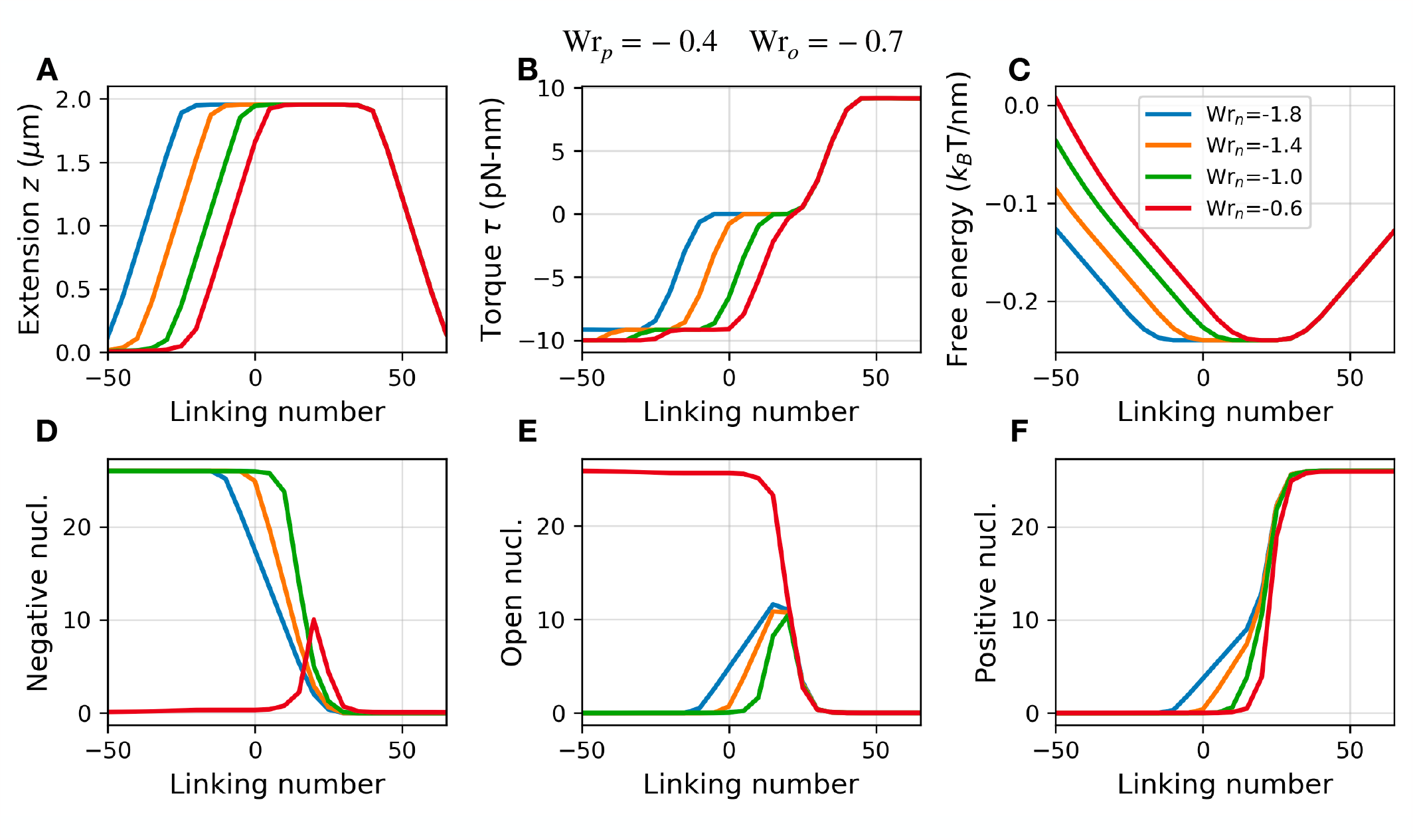
Torsional response of chromatin fiber with different writhe for negative nucleosome states. (A) Extension, (B) Torque, (C) Free energy, (D) Number of negative nucleosomes, (E) Number of open nucleosomes, and (F) Number of positive nucleosomes as a function of excess linking number being injected into the chromatin fiber. We used a reference of Wr_ref_ = *N* Wr_*n*_, corresponding to Fig. S1. Higher negative writhe increases the torque plateau as chiral transitions are able to absorb larger linking numbers, increasing the supercoiling range over which chiral states coexist. Same as seen previously (Fig. S6 and Fig. S5), when the writhe of the negative state is less negative than the open state (Wr_*n*_ =− 0.6), negative states are destabilized and the open state dominates the negative plectoneme regime. This, again, is a geometrically unphysical state if the nucleosome’s inner turn is not perturbed.

**FIG. S7.**
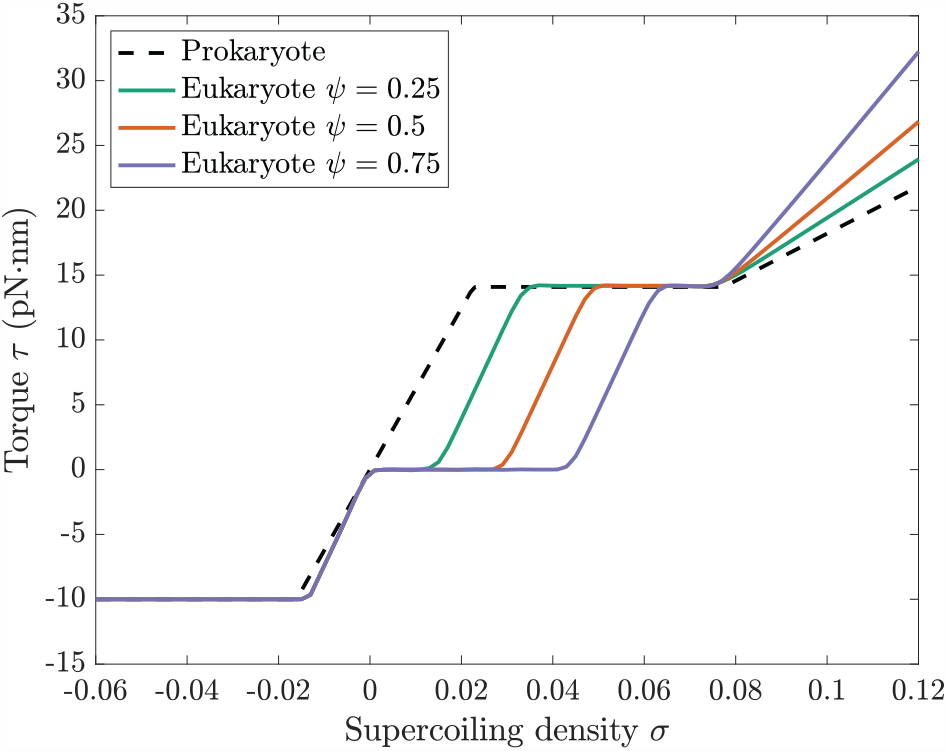
Comparison of the restoring torque for the case of prokaryotes (naked DNA) with the case of eukaryotes (DNA bound by nucleosomes). In the presence of nucleosomes, a plateau-like region emerges for *σ >* 0 wherein the restoring torque remains flat (*τ* ≈ 0) with an increase in *σ*. The size of this plateau region increases with an increase in the number of nucleosomes bound to the DNA segment (also shown in Fig. 1 D). In contrast, for the prokaryotic case with naked DNA, the restoring torque increases linearly as the DNA is twisted in the positive direction starting from *σ* = 0. Here, the behavior for a DNA / chromatin segment of length 10 kb under a force *f* = 1.0 pN is shown. The prokaryotic torsional response was calculated as described previously by Tripathi *et al*. [14].

**FIG. S8.**
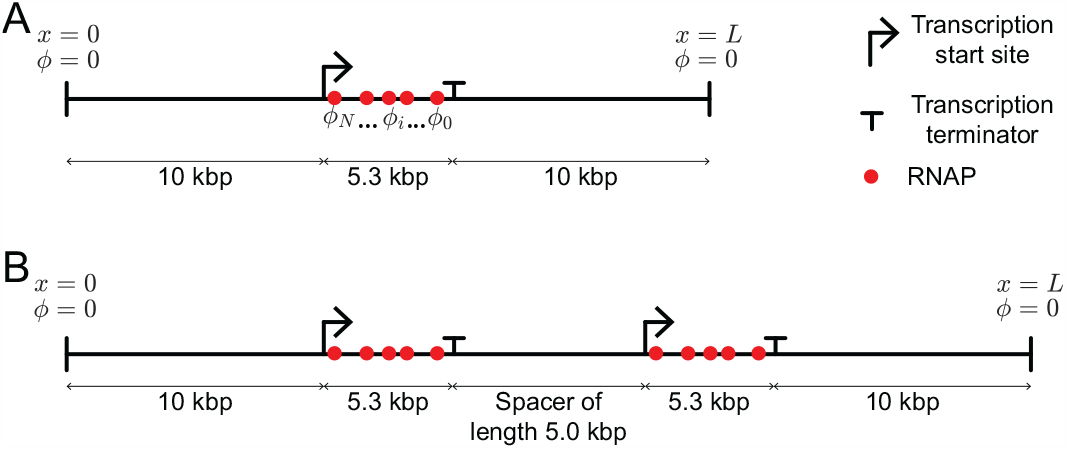
A schematic showing the different simulation setups used in the present study. **A** A single gene in a genomic segment with clamped ends (torsionally constrained DNA). This setup is used for the simulations in Fig. 2, 3, S9, S10, S11, and S12. **B** Two genes in a genomic segment with clamped ends. This setup is used for the simulations shown in Fig. S13. In each panel, *ϕ* indicates the DNA rotation angle at the RNAP sites or at the ends of the genomic segment.

**FIG. S9.**
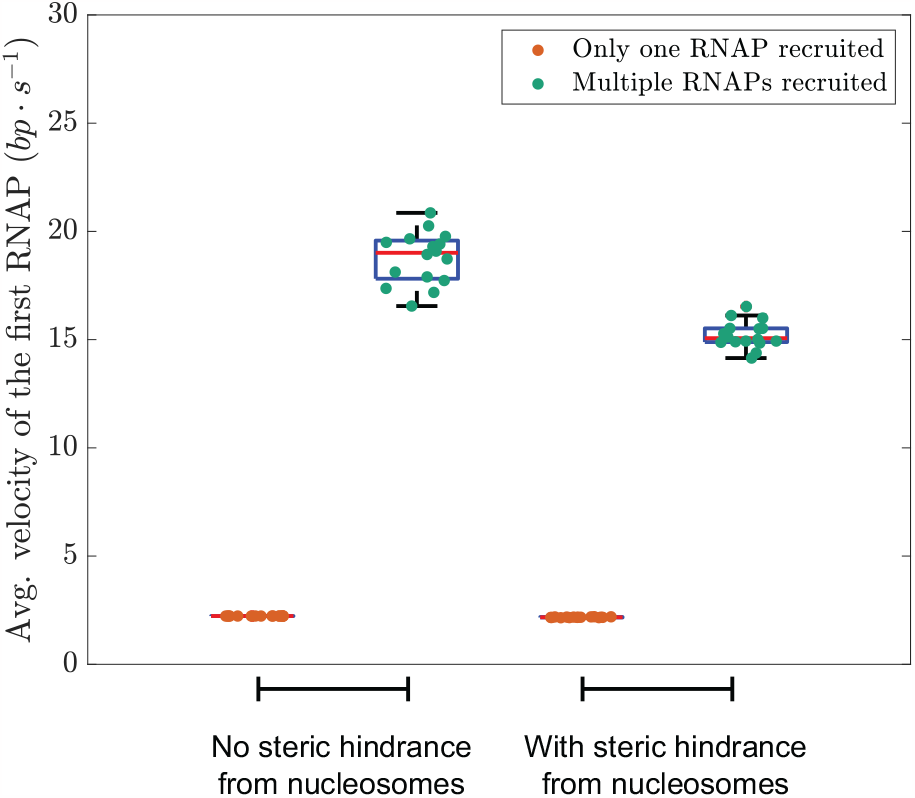
An RNAP moves faster if additional RNAPs are subsequently recruited to the same transcription start site. The figure shows the results for 16 independent runs with no supercoiling relaxation (*k*_*relax*_ = 0.0). For the cases with recruitment of multiple RNAPs, we used *k*_*on*_ = 0.5 min^−1^.

**FIG. S10.**
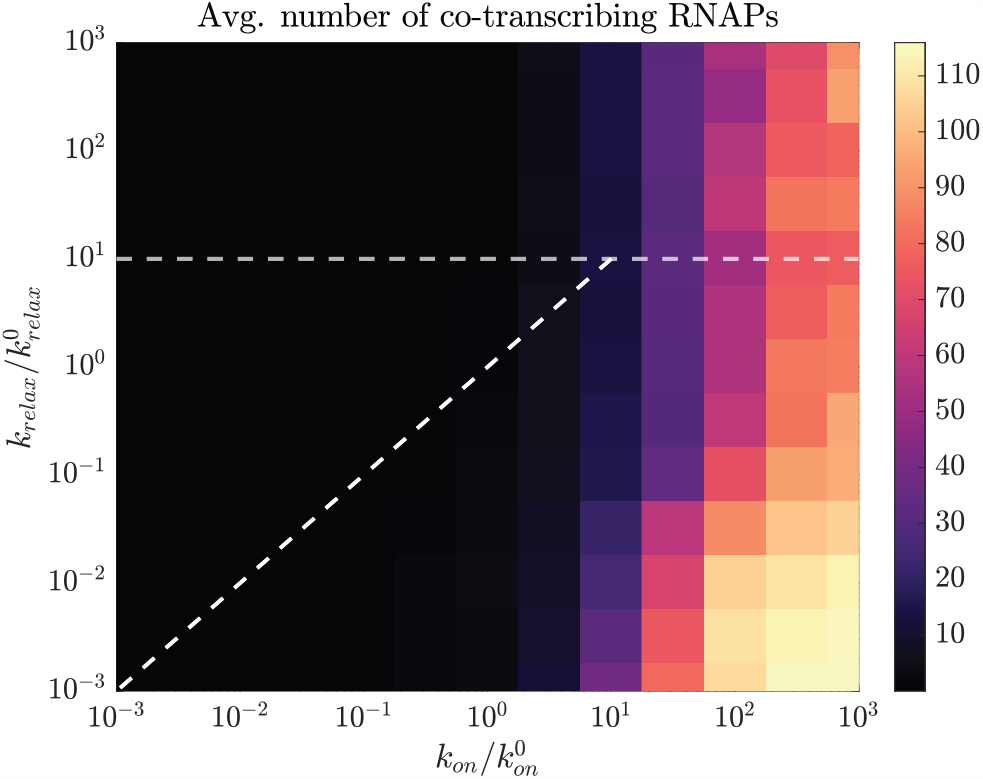
Average number of co-transcribing RNAPs during eukaryotic transcription for different values of *k*_*on*_ and *k*_*relax*_. Here, the simulation setup and parameters are the same as in Fig. 2 C; the dashed white lines correspond to the regimes shown in Fig. 2 C.

**FIG. S11.**
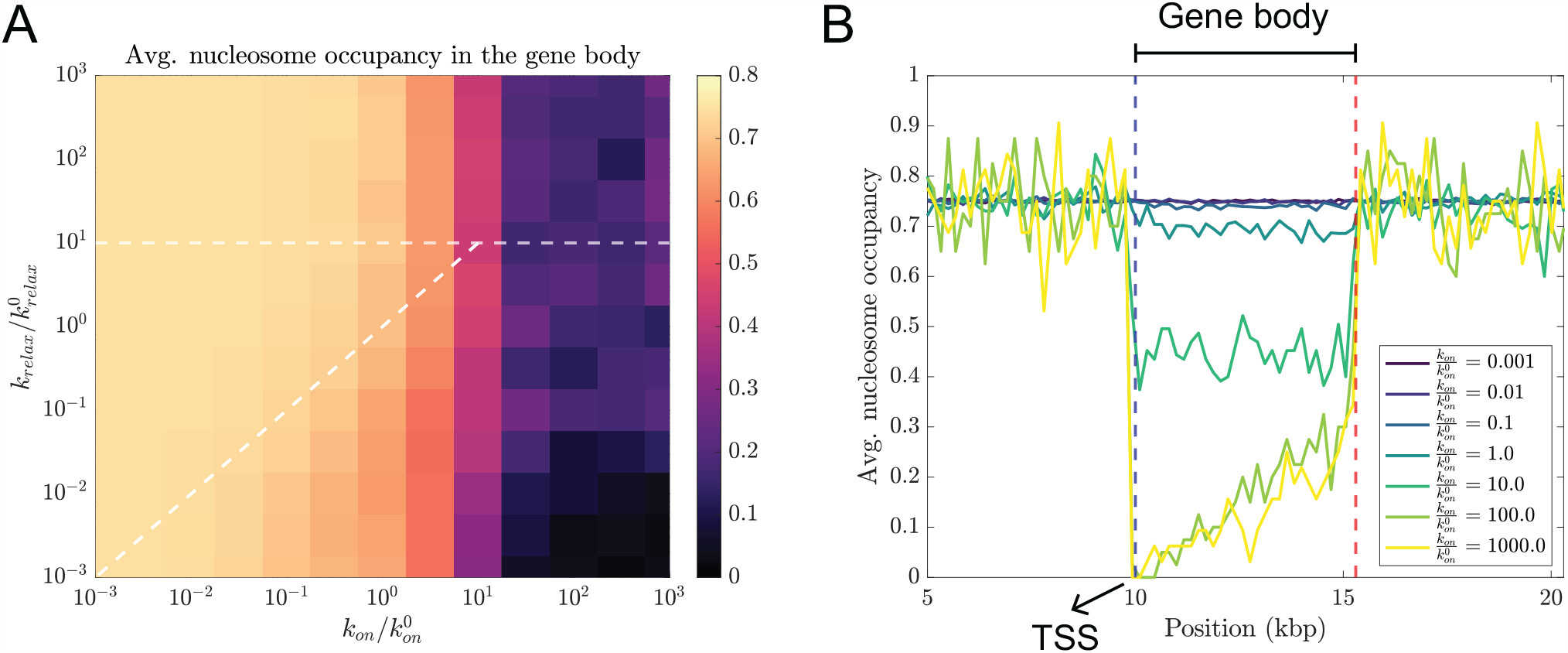
Nucleosome occupancy in the gene body can change in response to changes in the transcription kinetics. **A** Average nucleosome occupancy in the gene body for the simulation setup in Fig. 2 C. **B** Average nucleosome occupancy as a function of the distance from the transcription start site (TSS) for different values of *k*_*on*_. Nucleosomes are depleted from the gene body of highly transcribed genes due to the steric hindrance to nucleosome binding from RNAPs. The depletion is higher in the case of highly transcribed genes, and close to the TSS. In panel B, the simulation setup and parameters are the same as in panel A; panel B shows the behavior for a fixed value of 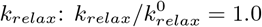.

**FIG. S12.**
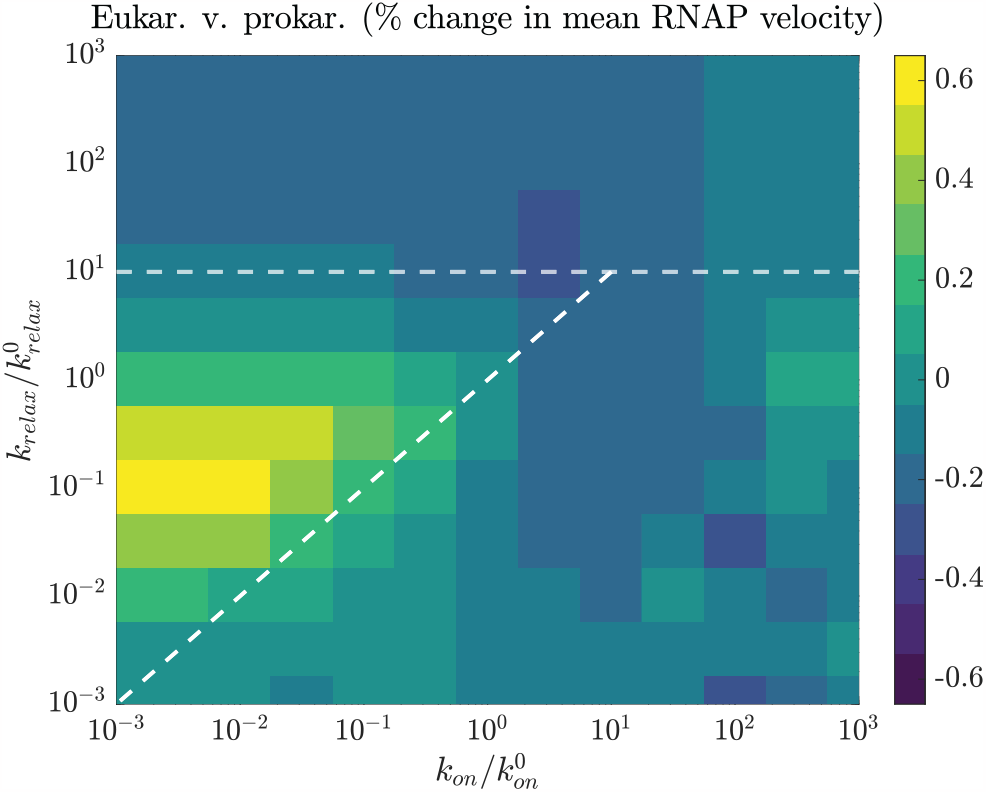
Comparison of the average RNAP velocity in prokaryotes and eukaryotes when nucleosomes in the eukaryotic genome offer steric hindrance to RNAP movement. The behavior is similar to the case wherein nucleosomes offer no steric hindrance (Fig. 2 C) and the dashed white lines in this figure are the same as in Fig. 2 C. Note that even the regime of fast supercoiling relaxation (high *k*_*relax*_; above the horizontal, dashed white line) and in the regime of supercoiling cancellation by co-transcribing RNAPs (below the inclined dashed, white line), the average RNAP velocity in eukaryotes is lower than the prokaryotic case: steric hindrance from nucleosomes is the dominating nucleosomal effect in these regimes with little contribution from nucleosomal torsional buffering (since supercoiling in these regimes is either quickly relaxed or cancelled). In the regime of low *k*_*on*_ and low *k*_*relax*_ (region enclosed between the two dashed, white lines), the torsional buffering effect still dominates, leading to a speed-up of transcriptional elongation in eukaryotes. Here, 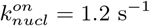 and 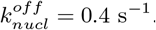.

**FIG. S13.**
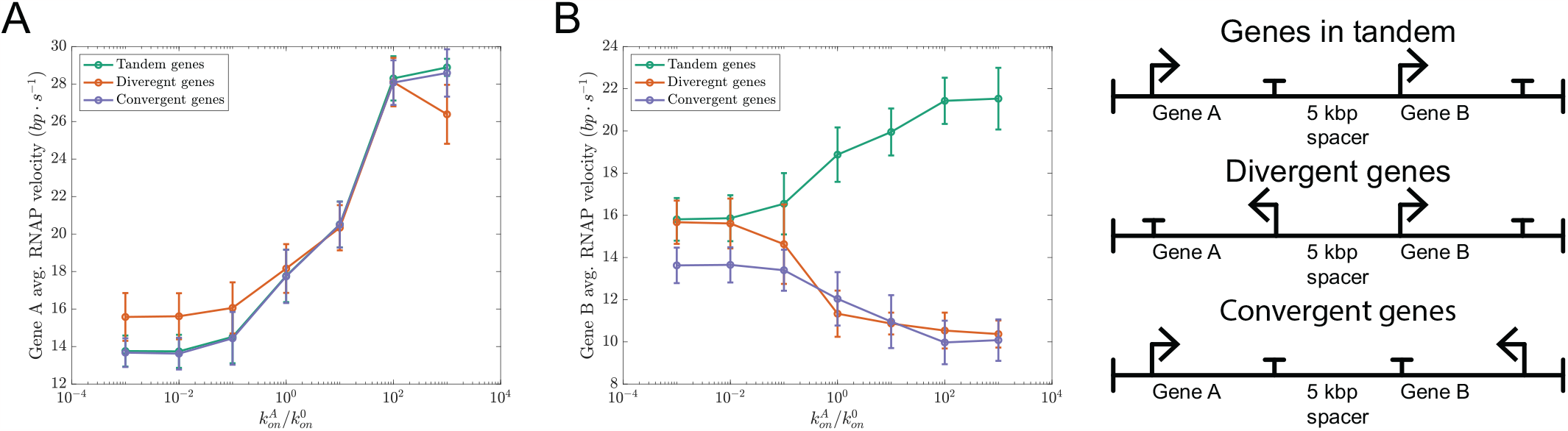
Gene orientation determines the nature of supercoiling-dependent coupling between neighboring genes. **A** Average RNAP velocities for gene A transcription when the transcription initiation rate for gene 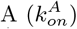 is varied. **B** Average RNAP velocities for gene B transcription when the transcription initiation rate for gene 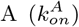 is varied while keeping the gene B transcription initiation rate 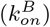 fixed. When gene A and gene B are oriented in tandem (see legend on the right), the average velocity of gene B RNAPs increases when gene A transcription is induced. This is because the negative supercoiling injected into the intergenic region by gene B transcription can be cancelled out by the positive supercoiling injected into the region during gene A transcription. When gene A and gene B are in divergent or convergent orientation, their transcription injects the same type of supercoiling into the intergenic region (negative for divergent genes and positive for convergent genes) resulting in supercoiling accumulation in the intergenic region. Consequently, in the case of both divergent and convergent genes, transcription of gene B slows down when gene A is highly induced. Here, 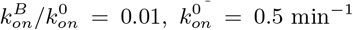, and *k*_*relax*_ = 5.0 min^−1^. Error bars indicate the standard deviation.

